# Temporal variability can promote migration between habitats

**DOI:** 10.1101/2023.04.23.537980

**Authors:** Harman Jaggi, David Steinsaltz, Shripad Tuljapurkar

## Abstract

Understanding the conditions that promote the evolution of migration is important in ecology and evolution. When environments are fixed and there is one most favorable site, migration to other sites lowers overall growth rate and is not favored. Here we ask, can environmental variability favor migration when there is one best site on average? Previous work suggests that the answer is yes, but a general and precise answer remained elusive. Here we establish new, rigorous inequalities to show (and use simulations to illustrate) how stochastic growth rate can increase with migration when fitness (dis)advantages fluctuate over time across sites. The effect of migration between sites on the overall stochastic growth rate depends on the difference in expected growth rates and the variance of the fluctuating difference in growth rates. When fluctuations (variance) are large, a population can benefit from bursts of higher growth in sites that are worse on average. Such bursts become more probable as the between-site variance increases. Our results apply to many (≥ 2) sites, and reveal an interplay between the length of paths between sites, the average differences in site-specific growth rates, and the size of fluctuations. Our findings have implications for evolutionary biology as they provide conditions for departure from the reduction principle, and for ecological dynamics: even when there are superior sites in a sea of poor habitats, variability and habitat quality across space determine the importance of migration.

## 1 Introduction

Migration is important to the life history of many species, with consequences for range shifts, genetic diversity, and population dynamics [49]. Traits governing migration are subject to selection and can significantly influence eco-evolutionary dynamics of spatially structured populations [16, 38, 9, 50]. In these papers, the term migration may mean ‘regular movements by animals between locations’ while dispersal may refer only to ‘movement of an individual from site of birth to site of reproduction’ and thus may be different behaviours [41, 11]. But we are concerned with a conceptually simpler problem: understanding the conditions that promote evolution of migration (here considered the same as dispersal) in the simplest sense of ‘moving from one spatial unit to another and back’ [6].

To explain why this is an important and interesting issue, consider what happens in the absence of variability, i.e., when population growth rate at each site is fixed. Qualitatively, for two sites, any dispersal from the better site (call it site 1) to the poorer site (call it site 2) would decrease the overall growth rate of the total population. Hence we would not expect dispersal away from the most favorable site to be favored by natural selection. The surprisingly complicated mathematical proof of this for multiple sites was given by [32]. Karlin proved that the overall population growth rate will be determined by the rate prevailing at the best site. Thus in constant environments, dispersal away from the best site can reduce population growth. That has since become established as one example of a general reduction principle which we may describe as ‘mixing reduces growth’ and ‘differential mixing selects for reduced growth’ in constant environments [7, 29, 23, 1, 3, 2, 4].

Past work suggests that environmental variability may select for dispersal, as it allows populations to cope with variation in resources, escape unfavorable environments and benefit from new environments [53, 8, 5, 52], thus providing conditions for a departure from the reduction principle [42, 2]. Environmental or resource variability has been incorporated by accounting for temporal variation in carrying capacities [39, 15] or growth rates [63, 34], using approximations and/or numerical simulations to examine the evolution of dispersal. Many other studies have also shown that dispersal is relevant in spatially and temporally varying environments [62, 24, 31, 15, 43, 56, 42, 51, 30, 46, 40].

In this paper we answer a simple question: are there conditions that would favor the evolution of dispersal among sites with distinct stochastic growth rates? The use of stochastic growth rate as the appropriate invasion fitness into a monomorphic population has been established in both a genetic context [59] and a general ecological context [44]. We are interested in migration evolution and invasion dynamics, and therefore do not consider the extensive work on how population density affects dispersal [57].

The initial condition for an evolutionary analysis is the case where there is no dispersal. In the neighborhood of that initial condition, a central challenge for this analysis is that the stochastic growth rate changes very rapidly and is non-differentiable (a singularity) in the case of no dispersal. What does this mean? Let *m* be the probability of migration between two sites. The system of matrices will not satisfy at *m* = 0 the conditions under which the population growth rate is known [64] to be analytic. Our results show that under some conditions (as we describe further below) there is indeed a singularity at *m* = 0, with the derivative going to ∞. Thus, a typical approach such as a perturbation analysis cannot predict the exact nature of migration rate near *m* = 0. This can be seen in simple numerical simulations of migration between two sites. Such a singularity has been pointed out in special models for plants [48], life cycle delays [61], dispersal [63] and structured plant models [10]. Note that kin selection and density-dependence are not able to alter this singularity. We address the challenge by establishing rigorous inequalities for stochastic growth rate.

To describe the problem and our model, we first consider the simple case of two sites with density-independent exponential population growth rates. We assume that population growth rates change over time and the logarithm of growth rates is normally distributed. We characterize the stochasticity of environmental conditions by the variance (*σ*^2^) of difference in site-specific population log growth rates (allowing for covariation across sites). Migration is parameterized by a probability *m* of changing site and (1 − *m*) of staying.

We assume there is a superior site (site 1), meaning that its expected logarithmic growth rate µ_1_ is greater than the expected logarithmic growth rate µ_2_ at site 2. Thus, populations have to overcome an average growth rate difference between sites, given by 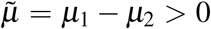. Our metric for fitness is the (long-run) stochastic population growth rate, *a* [58]. We examine the asymptotic limit, (*i*.*e*., small migration limit) of the long-run population growth rate *a*(*m*).

We find the change *a*(*m*) − *a*(0) in the stochastic population growth rate with migration relative to no migration is bounded above and below by functions that behave as a power *m*^*γ*^ of the migration rate. When *γ <* 1 there is, as mentioned earlier, a singularity at 0: the derivative of *m*^*γ*^ is proportional to (1*/m*^1−*γ*^) (which blows up as *m* approaches 0). Thus, when *γ <* 1 the function *m*^*γ*^ is non-differentiable at 0, and will dominate any linear function for *m* sufficiently close to 0. Under general assumptions we prove that the power *γ* of the migration rate is bounded above by 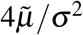. Therefore, small migration is favored relative to zero migration when the variance of difference of growth rates between sites is large enough to over-come a fixed difference in expected growth rates. The intuition is that when there are independent fluctuations in growth rates across sites, populations may use dispersal to exploit this environmental variability (*σ*^2^) to their benefit. As the difference 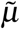 in average quality increases, even larger fluctuations are required to make migration favorable.

Note that these results refer only to the variance of the difference in log growth rates between the sites. Whether these rates are positively correlated, negatively correlated, or uncorrelated depends in addition on the variance at each site individually. These individual variances will affect the baseline growth rate *a*(0), but have no bearing on the results we discuss here concerning the difference *a*(*m*) − *a*(0). In addition, we show that migration can be advantageous in spite of costs proportional to the total migration rate — if, for instance, we suppose that some proportion of those migrating are lost or fail to reproduce. We consider as well the setting where there are multiple sites, where it may not be possible to migrate in one step between the best and second-best sites. We consider the impact of path lengths on the long-term growth rate during migration and find that short paths to return to the best site are favored. We also find that path lengths matter only when sites are similar (on average).

The paper provides an approach to studying migration patterns (such as those of birds or mammals) by examining variation in environmental conditions, and describing the relationship with site-specific metrics (such as population growth rates), to help bridge the gap between theory and empirical studies [40]. The next section of the paper describes the model in constant and variable environments. We follow with the theorem and simulations for the two-site case. Next, we describe the general theorem for any number of sites and simulations for 4 sites. We conclude with further discussion of the relevance of our results.

## 2 Model in constant environments

We describe the model by first considering constant environments, and then introduce variable environments. The growth rates of the population at each site are described by a diagonal matrix **G**, where each diagonal element represents site-specific growth rate. There is one “best” site at which the growth rate is higher than at every other site. We assume no individuals die or reproduce during migration, and dispersal matrix **M** is stochastic (the rows are probabilities of moving from or staying at a site and must sum to 1). The individuals can get from the one site to any other site in a finite number of steps, so the migration matrix **M** is irreducible. (Throughout bold symbols refer to matrices and vectors.)

Let the components of vector **N**(*t*) = (*N*_1_(*t*), *N*_2_(*t*))^T^ denote the population size at each site and T denotes the transpose. Then the population dynamics can be described by

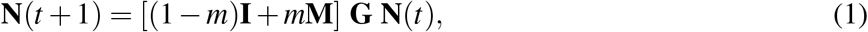

where the migration rate *m* scales the pattern of dispersal described by **M**.

When there is migration, that is *m*≠ 0, the overall growth rate is given by the largest eigenvalue (also called spectral radius **s**) of the product of **M**(*m*) = [(1 − *m*)**I** + *m***M**] and diagonal growth matrix **G**. [32] used the Donsker–Varadhan formula for principal eigenvalues to prove that the spectral radius is strictly decreasing in migration rate *m*, so in this case migration will not favored. We would like to note that Karlin’s Theorem 5.2 was independently derived by [35], using a very different proof (though still deeply connected to Donsker–Varadhan theory).

Interpreted biologically, the Karlin’s theorem states: in such a system, the greater the level of mixing of population between sites, the lower the long-term growth rate of the total population. Such selection for reduced dispersal under constant selection is called the *reduction principle*, studied primarily in a genetic context by [21, 22, 23]. The reduction principle has since been extended to the evolution of mutation rates, sexual reproduction, and cultural transmission [3, 2].

### Incorporating time-varying growth rates

Thus dispersal is not favored in a model with fixed growth rates. Now assume population growth depends on site-specific environmental factors that change over time. To account for this variation we treat growth rates as a random process and equation (1) is modified using a time-dependent matrix specifying growth-rates (**G**_**t**_) as:

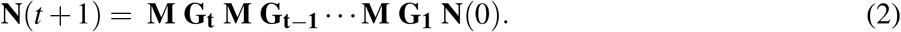

Analysis of such a system is challenging for many reasons [17, 48, 2]. For structured populations, the stochastic growth rate is notoriously difficult to compute. Thus, we look beyond the domain of classical probability to the theory of large deviations. That theory examines extreme events that emerge from multiple rare occurrences. In our setting, the rare migration events can accumulate over time to dominate the aggregate dynamics, as described in the model and analysis below.

## 3 Model for heterogeneous environments

To analyze dispersal evolution, we assume density independence, and for simplicity assume no age or stage structure (i.e., a scalar population) at each site. We start with two sites (1 and 2), where population grows at a rate 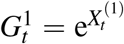 and 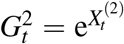, respectively. The log growth rate vector 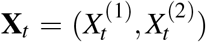 at *t* is assumed to be serially independent; however, the components within each **X**_*t*_, corresponding to growth rates at different sites, may be dependent.

The expected values of log growth rates are given by 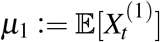 and 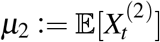 for site 1 and 2, respectively. We assume site 1 is the best site on average, meaning µ_1_ is strictly larger µ_2_. So the difference 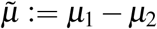 between average growth rates is always positive 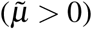.

We assume the difference in growth rates between site 1 and site 2 at each time *t* is distributed normally, and 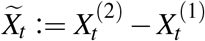 has variance *σ*^2^. Note that *σ >* 0 so long as the two sites are not perfectly positively correlated. This allows us to vary site-specific variances and co-variances between sites. The arguments and theorems hold whether growth rates 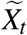 are assumed to be Gaussian or sub-Gaussian. Sub-Gaussian refers to a class of random variables whose tails decay at least as fast as a Gaussian (normal) distribution, meaning that extreme values are no more likely than for a Gaussian variable. This means that our results are not tied to the details of the Gaussian distribution, but only to the general shape of the extreme-event likelihood. At the same time, while the results do still hold when tails are strictly **thinner** than Gaussian tails — for instance, when the growth-rate fluctuations are bounded — the result is much less interesting in this case, as the both bounds are then strictly negative for small *m*.

The probabilities of migration to another site and of staying at a site are given by *m* and (1 − *m*), respectively. The populations at the two sites are governed by the system of equations

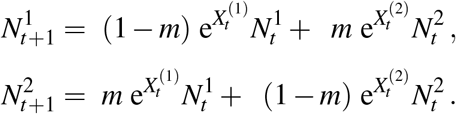

In matrix form the equations are written as

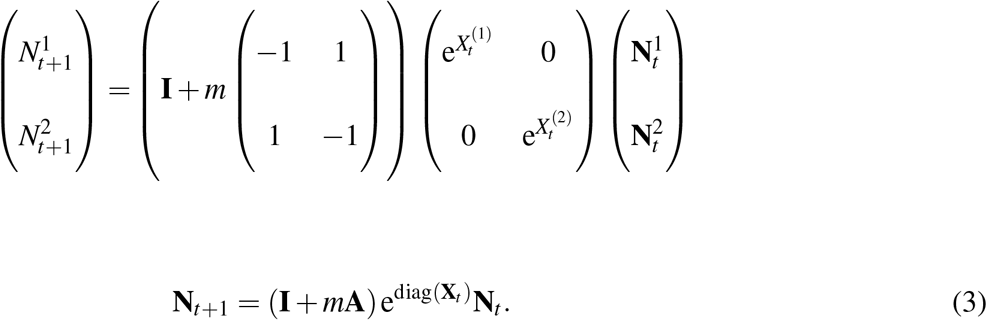

In the last equation we define 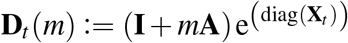, where the matrix **X**_*t*_ is diagonal with elements being the population growth rates 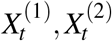 at the two sites. We assume that the sequence **X**_*t*_, **X**_*t*−1_, …, **X**_1_ is i.i.d. in time, although there can be any correlation between sites at the same time. Also, the pattern (but −1 1not the scale *m*) of dispersal is given by the matrix 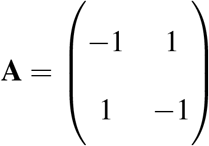. In the main manuscript, we assume the dispersal matrix **A** to be constant but the appendix generalizes the results to **A**_*t*_ matrices that are i.i.d with time.

### Long-run stochastic growth rate

Owing to the multiplicative nature of population growth (survival and reproduction), in a stochastic environment, the long-term growth converges to a limit as:

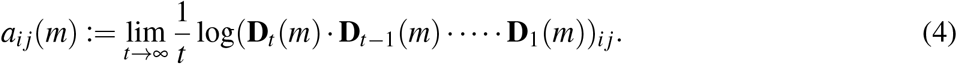

For *m >* 0 the i.i.d. sequence 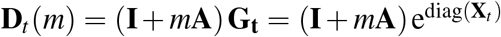 satisfies the conditions for the existence of a stochastic growth rate independent of starting condition [17] and so *a*_*i j*_(*m*) is the same for all *i* and *j*; we call this value *a*(*m*). When *m* = 0, the limit *a*(0)_*i j*_ in (4) exists, but is equal to µ_*i*_ at place *i* on the diagonal and −∞ off the diagonal. Thus, when there is no migration site 1 dominates the total population.

## 4 Main result for two sites in variable environments

### 4.1 Asymptotic result for two sites

For migration to be favored, the stochastic growth rate *a*(*m*) must increase as a function of migration rate relative to no migration, *a*(0). Our theorem below describes lower and upper bounds for *a*(*m*) − *a*(0) in the small-*m* limit in terms of the expected growth rates between two sites, 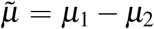 and the variance *σ*^2^ describing the fluctuations.

The probabilistic reasoning for how the theorem works is as follows: since the environment is fluctuating, there is always a fraction of times when the site with lower growth rate is doing better than the best site. The proof enumerates sequences of possible excursions (and their probabilities) for which the less favorable site does better than the favorable site. Here, excursions are defined as the set of paths in the migration graph that start and end at the best site 1, with no intervening returns to site 1. We would like to note that for the two-site case, the only possibility is to move from site 1 to site 2 and back. Excursions become more complex when we discuss the case for more than two sites in section 5. Once the excursions are defined we are able to balance the exponentially growing number of excursions against the exponentially large probability penalty incurred by repeated migration events to find a growth rate that will be achieved with probability 1.

Note that to ease the exposition the results for the case of two sites and more than two sites are presented separately in the main manuscript. In the Appendix, we present the general theorem for any number of sites. The mathematical details of the proof are presented in Appendix B.

#### Theorem 1.

*The difference between stochastic growth rates, a*(*m*) *and a*(0) *is bounded below and above (scaled by a constant) as follows: For any positive* δ,

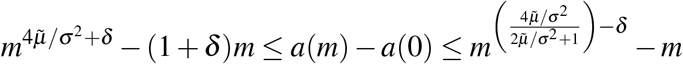

*for positive m sufficiently close to 0*.

Here *m* is the probability of migration, 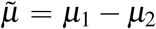 is the difference in expected growth rates between site 1 and site 2, 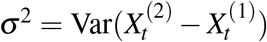 is the variance in difference between growth rates at site 1 and site 2 (at time *t*), *a*(*m*) is the stochastic growth rate with migration, *a*(0) is the stochastic growth rate without migration, and δ *>* 0 is a small positive quantity.

Note that the upper bound is dominated by the lower power of *m*, and the lower bound is dominated by the higher power of *m*. We are interested in the lower bound 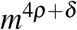 for small *m*, where 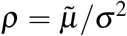. Note that the increase in growth rates is at least as large as 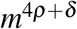. The smaller the ρ, the more steeply the power 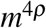 jumps up from *m* = 0. For the ratio ρ to be small either the numerator (the expected inter-site difference 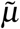) must be small or the denominator (the variance of inter-site difference *σ*^2^) must be large. To put it differently, the evolution of migration will be favored when the variance of inter-site log growth rate difference (*σ*^2^) is large enough to overcome a fixed mean difference 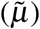. In the Appendix A, we present a figure (Figure 2) and a table with upper and lower powers, 4ρ and 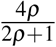, respectively. We simulate the slope as 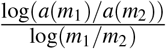 by taking two values of migration rate (*m*_1_ = 1 × 10^−12^ and *m*_2_ = 1 × 10^−10^). Most of the simulated slopes lie within lower and upper powers as shown in Figure 2 in the Appendix.

**Figure 1.**
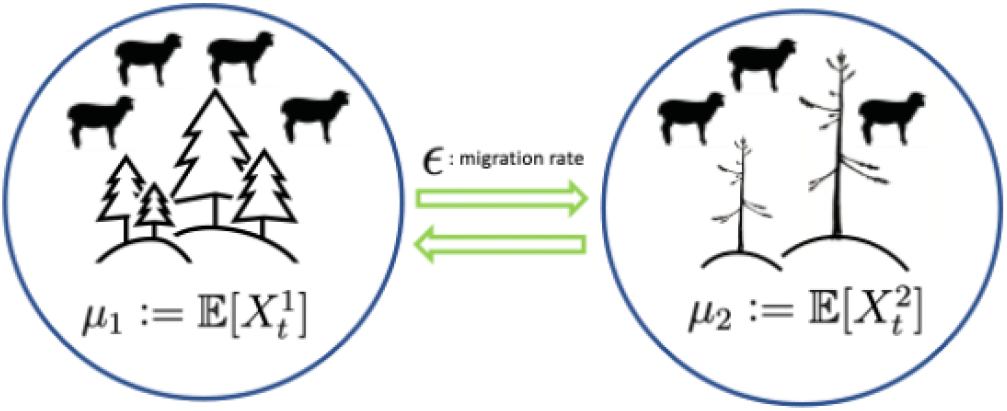
Site 1 (best site) has higher expected growth rate (µ_1_) than at site 2 (µ_2_).

**Figure 2.**
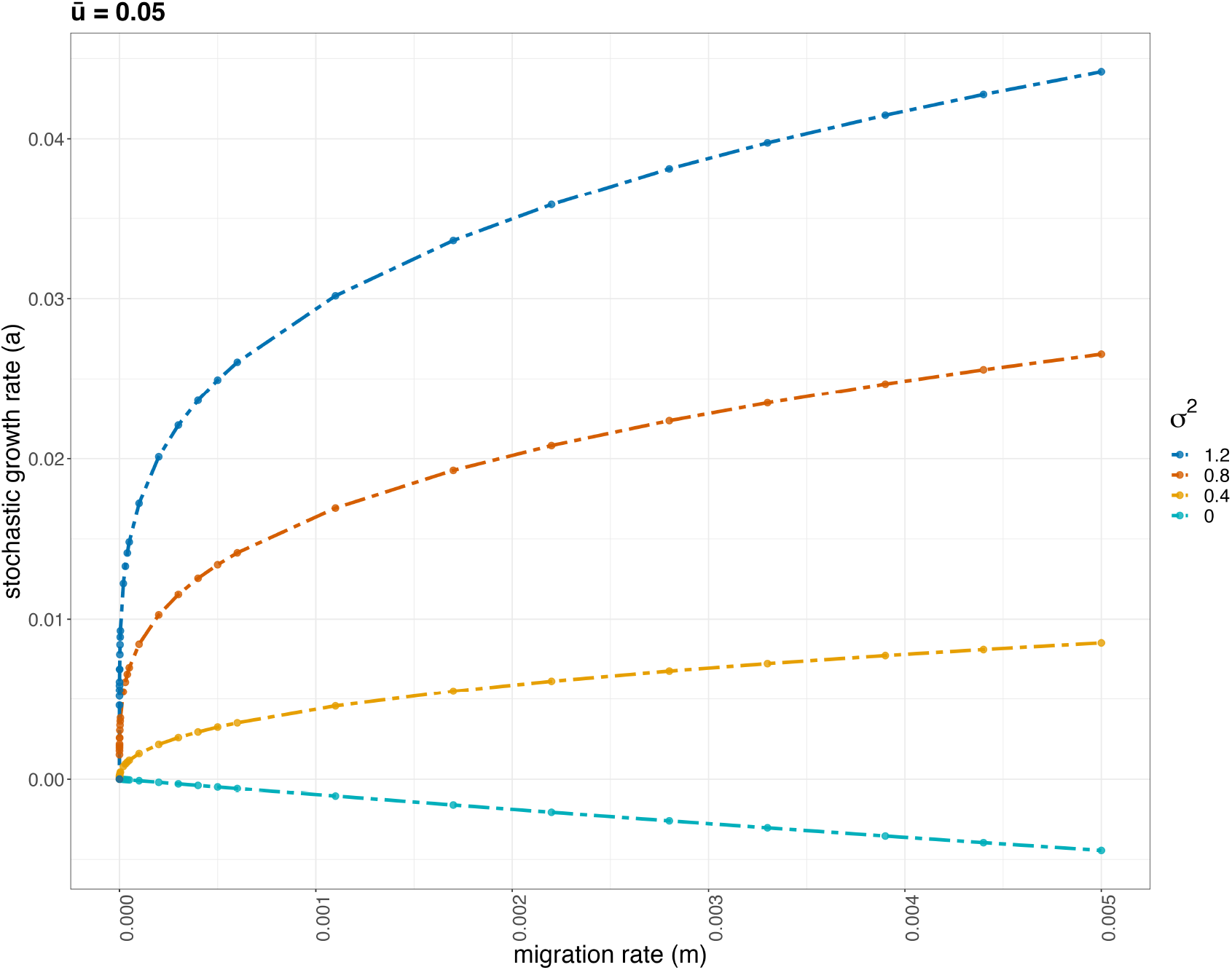
Stochastic growth rate as a function of migration rate. Different colors indicate different values of variability, described by *σ*^2^. Higher values of *σ*^2^ correspond to increase in stochastic growth rate for a fixed value of difference between average site-specific growth rates 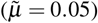. Here, *a*(0) is 0.

Note that this increase 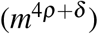 is opposed by a linear penalty −(|**A**(0, 0)| + δ)*m*, corresponding to the direct effect of population lost to the steady stream of out-migration from the optimal site. The linear penalty is of the form |**A**(0, 0)| since δ can be made arbitrarily small. Since the dispersal matrix is constant **A**,−(|**A**(0, 0)| + δ) is just (1 + δ). In the Appendix B, **A** can vary with time (**A**_*t*_) and hence the linear term is of the form of expectation of absolute value of **A**_*t*_(0, 0). When ρ is smaller than 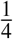, the positive term will definitely overwhelm the linear penalty for sufficiently small positive values of *m*. For large values of ρ the linear migration penalty will definitely predominate, and no positive amount of migration will be favorable. The theorem does not exclude the possibility that this transition may occur at an even higher level, up to 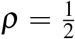 (when the power on the right-hand side is 1). But once the power of *m* is known to be below 1, this migration benefit persists for sufficiently small *m* regardless of any additional migration costs proportional to *m*.

In the next section, we use simulations to illustrate and examine the effects of 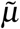 and *σ*^2^ through simulations. Following that, we use our simulation results to present an intuitive argument (section 4.3) for our analysis.

### 4.2 Simulations for two sites

We illustrate the theorem by examining the effects of varying fluctuations (*σ*^2^), the difference in average growth rates between sites 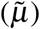, or both on stochastic growth rate *a*(*m*). The stochastic growth rate can be calculated from numerical simulations of many time steps of a population.

#### 4.2.1 Effect of varying fluctuations on stochastic growth rate

We first find that when the difference in mean growth rates between the sites is small (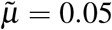 in Figure 2), a small positive migration rate will benefit from the variability between sites (described by *σ*^2^) relative to no migration. As predicted by the theorem, increasing the variability *σ*^2^ of the difference between 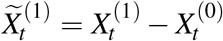 increases the benefit from migration. This may enable populations to grow faster than would have been possible at the single best site.

Our finding suggests that a species will benefit from the ability to wander between two similarly good territories, where environmental conditions vary separately between them from year to year. We discuss the intuition for the ratio 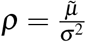 at the end of the section.

Figure 2 describes stochastic growth rate *a* as a function of migration rate *m* for different levels of variability, characterized by *σ*^2^. For low values of *σ*^2^, the gain from migration is not sufficient to cause an overall increase in the long-term growth rate. However, for higher values of *σ*^2^, introduction of even small level of migration leads to a sharp increase in stochastic growth rate. The sharp increase with migration (neighborhood of *m* = 0) is evident in Figure 2 and in the log plot as shown in Appendix A. Therefore, given enough variability (a high enough value of *σ*^2^), migration rate *m* can increase *a*(*m*) relative to *a*(0).

#### 4.2.2. Effect of varying difference in mean growth rates between sites

Next we consider the effect of changing the difference 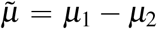 between the average growth rates at sites 1 and 2. We find that the larger the difference between the expected growth rate in sites (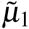 increased from 0 to 0.45 in Figure 3), the smaller the benefit from migration, in terms of increase in long-term growth rate.

**Figure 3.**
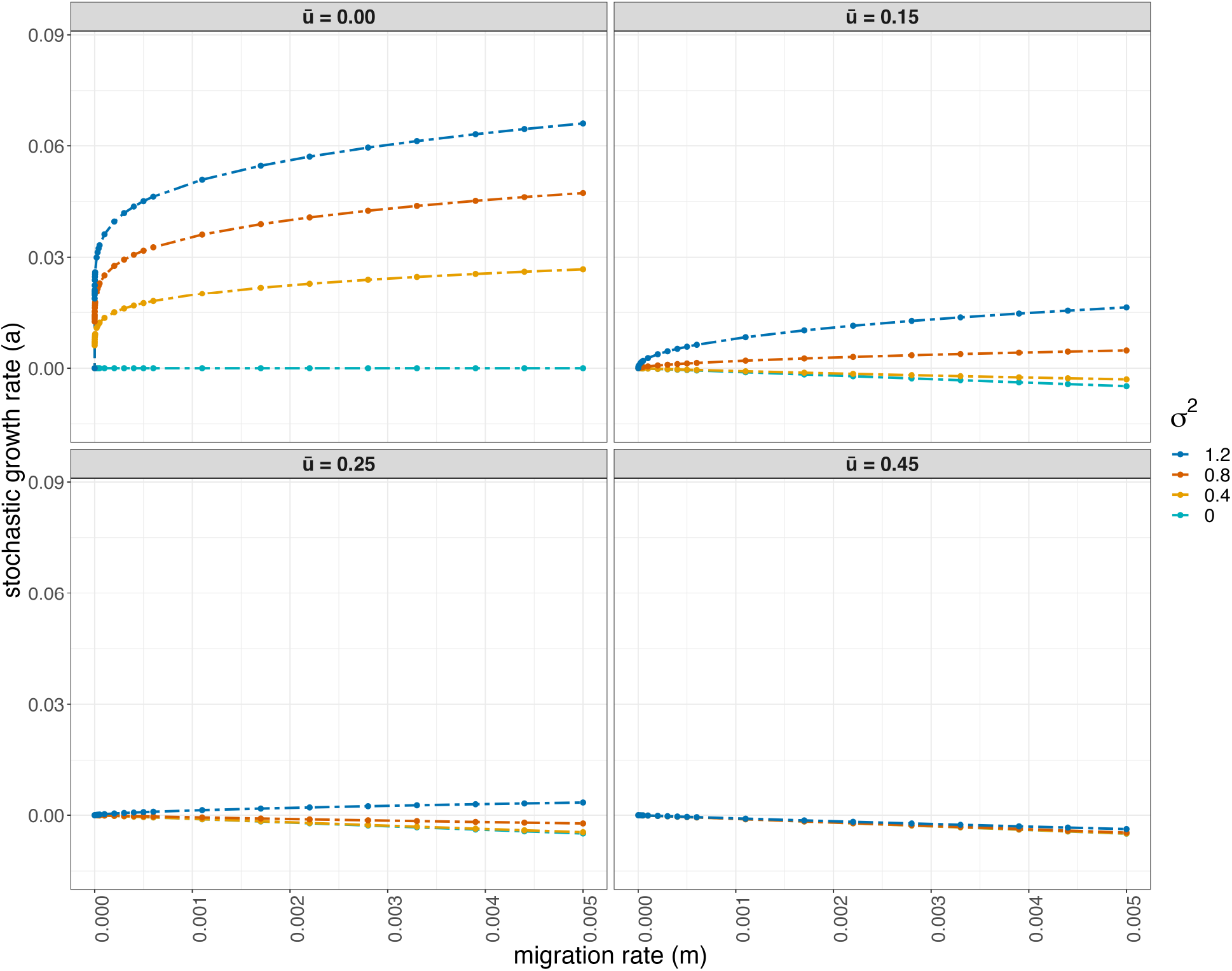
Stochastic growth rate as a function of migration rate. On the x-axis is the migration rate, on the y-axis is the stochastic growth rate. Each panel corresponds to a fixed value of difference in expected growth rates between two sites, 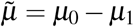. Different colors indicate values of variability, *σ*^2^.

Biologically, a greater difference 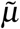 will make the rare events when growth at the inferior site exceeds the growth at the best site even rarer, while a greater variance (*σ*^2^) will make such exceptional deviations more likely. Thus, high variance may compensate for a large difference in mean growth rate but only to an extent. The ratio determines whether populations may be better off without migration.

Figure 3 depicts the effect of changing 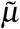 on stochastic log growth rate *a* as a function of *m*. (The growth rate at site 1 is set to 0.) The first panel (top-left) presents the case where both sites have same expected growth rate 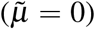. In this case, when variability *σ*^2^ equal to 0 the stochastic log growth rate is always 0, the common growth rate of the two sites (light blue line). For any nonzero *σ*^2^, *a* jumps up rapidly as *m* departs from 0. (Note that the case 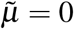 is not actually covered by the results of this paper. It is shown in [54] the increase in *a*(*m*) as *m* increases from 0 is faster than any power law.) In the last panel, by contrast, 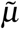 is so large that the stochastic growth rate does not increase for any of the given fluctuation sizes. There is no benefit from migration, as the sites differ too greatly in their mean quality.

We illustrate the combined effect of changing site quality 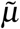 and variability *σ*^2^ on the difference in stochastic rates (with and without migration) *a*(*m*) − *a*(0) in Figures 4 (a) and (b). To emphasize the role of fluctuations in increasing the difference *a*(*m*) − *a*(0) (denoted by “a_diff” in the legend in Figure 4 (a)), the limits on the x-axis are above 0.5. In Figure 4 (a) it is evident from the contours that for a fixed difference in expected growth rates between sites 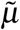, with enough variability *σ*^2^, populations overcome the effect of migrating to a site with poorer habitat quality. In Figure 4 (b), the blue and orange color contrast the gain in growth rate from migration due to higher *σ*^2^.

**Figure 4.**
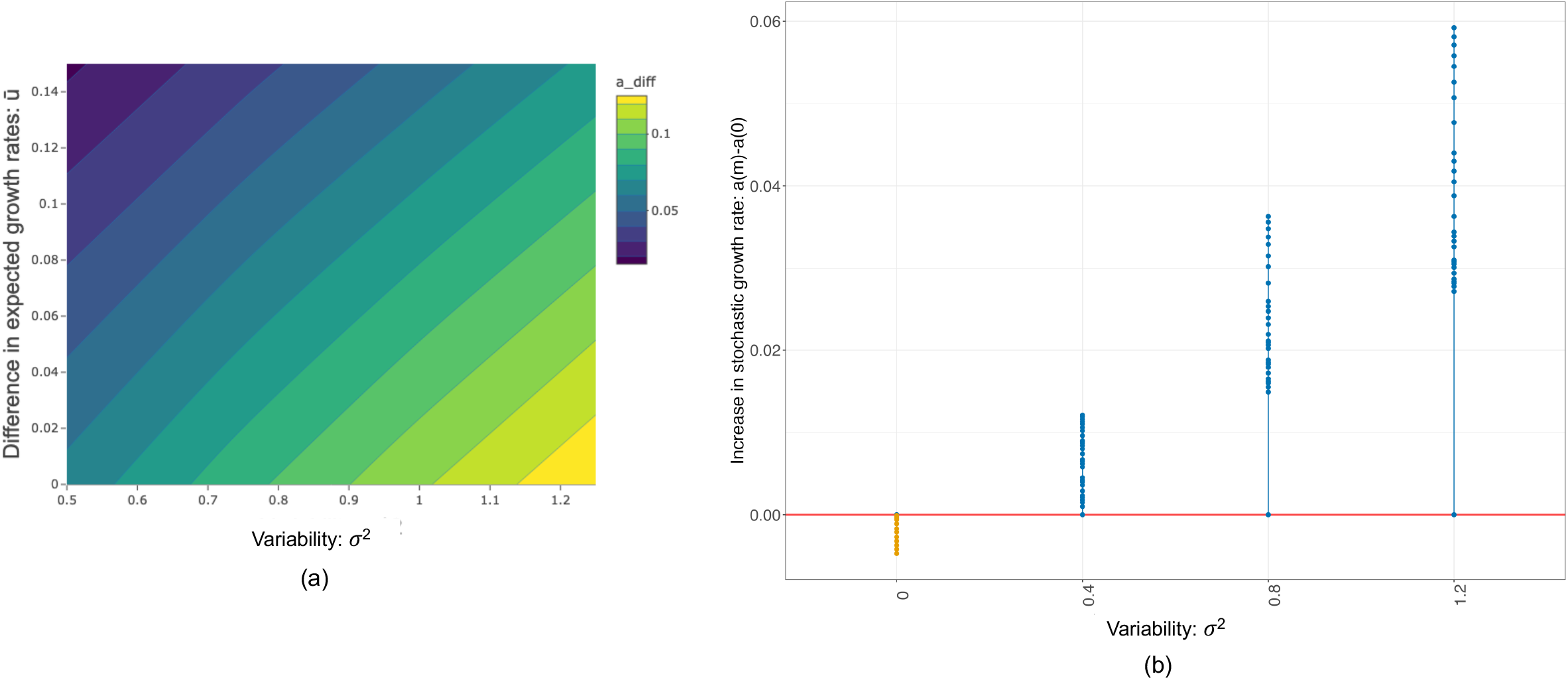
(a) The plot depicts the combined effect of 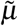 and *σ*^2^ on stochastic growth rate. On the y-axis is the difference in expected growth rates 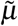. On the x-axis is the variability, *σ*^2^. Each color in the legend (“a_diff”) corresponds to values of *a*(*m*) − *a*(0) (we fix a value of migration rate *m* at 0.005). (b) Here, on the y-axis is difference in stochastic growth rate, with and without migration, *a*(*m*) − *a*(0). On the x-axis is the variability, *σ*^2^ at the fixed value of 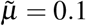 On the x-axis is the variability *σ*^2^. The blue and orange color is to contrast the gain in growth rate from migration due to higher *σ*^2^.

### 4.3 Heuristic arguments for the two-site case

Theorem 1 gives the lower and upper bound for the difference in stochastic growth rates (with and without migration), *a*(*m*) − *a*(0). Since the lower bound is positive and increasing as a function of *m*, migration increases the overall growth rate on the long run. The bounds depend on the ratio 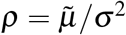. We discuss an intuition for the ratio. A similar ratio is discussed in [39] based on the mean and variance of the lognormal distribution.

In the case of no stochasticity, *X* ^(1)^ *> X* ^(2)^, and dispersal should not be favored as the population achieves a higher growth rate in site 1. When some individuals disperse during a time interval from site 1 to site 2, the disperser’s growth rate relative to a non-disperser is 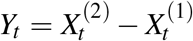. The geometric growth rate, per unit time, is changed by a factor exp(*Y*_*t*_). For *m* = 0, the expected growth rate, 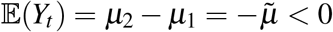 and Var(*Y*_*t*_) = *σ*^2^ *>* 0. Since we are assuming 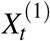 and 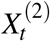 are normal, the distribution of *Y*_*t*_ is lognormal and 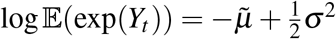. So the expected relative growth rate is greater than 1 if the ratio 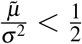.

This is one approach to understand why the bounds depend on ratio of 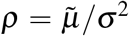, and also why 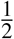 is an important threshold. However, the actual relative rate depends on the time period *T* over which a migrating ancestry experiences this relative rate, a concept that is formalised in section as the duration of an excursion (see Appendix B).

#### 4.3.1 Effect of varying migration penalty

Dispersal may be expensive if dispersing individuals are vulnerable to incremental risks of death [16, 9]. Therefore, we examined the effect of increasing migration penalty or costs on the long-run growth rate. The costs are increased by changing the elements in the migration matrix to account for individuals lost during migration. We make the row sum ∑_*i*_ **A**(*j, i*) negative to account for loss of individuals as individuals leave the best site at the rate −*m*.

This imposes an additional linear cost (as a function of migration rate *m*) to migration, which may make itself felt when *m* is large. Recall the lower bound in Theorem 1 is 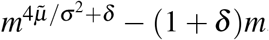. For small *m* the balance of costs and benefits for migration at rate *m* depends essentially on 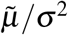. When this exponent 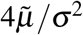 is smaller than 1 the benefit will dominate and growth will be increased for sufficiently small positive *m*, regardless of the size of the linear migration penalty. That is, assuming the costs of migration (borne by the individuals migrating) are linear in *m*, the advantage of small migration will persist even when migration is inherently costly. When the power of *m* is greater than 1 the migration penalty will dominate for small *m*, and the effect of migration on log growth rate will be negative.

The important point regarding introducing a cost to migration is that the effect in **A** is always nearly linear in *m*, while the increase of *a* near 0 is often superlinear, growing as *m*^*γ*^, where 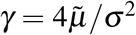. If the power *γ* is strictly less than 1, the rapid increase in *a* near 0 will be qualitatively unaffected by a linear term for *m* sufficiently small.

Figure 5 shows two panels with increasing levels of migration penalty at fixed 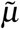 and *σ*^2^. The two plots in the left and right panel of Figure 5 correspond to high and low variability (*σ*^2^ = 1 and *σ*^2^ = 0.25). The penalty is introduced by making ∑_*i*_ **A**(*j, i*) negative by 20% and 10% as this implies individuals were lost during migration. In the top panel, we apply a migration penalty of 20% for a fixed 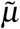 whereas the bottom panel corresponds to migration penalty of 10%. Despite a higher penalty in the top panel, we see that benefit from migration is attained when *σ*^2^ is higher.

**Figure 5.**
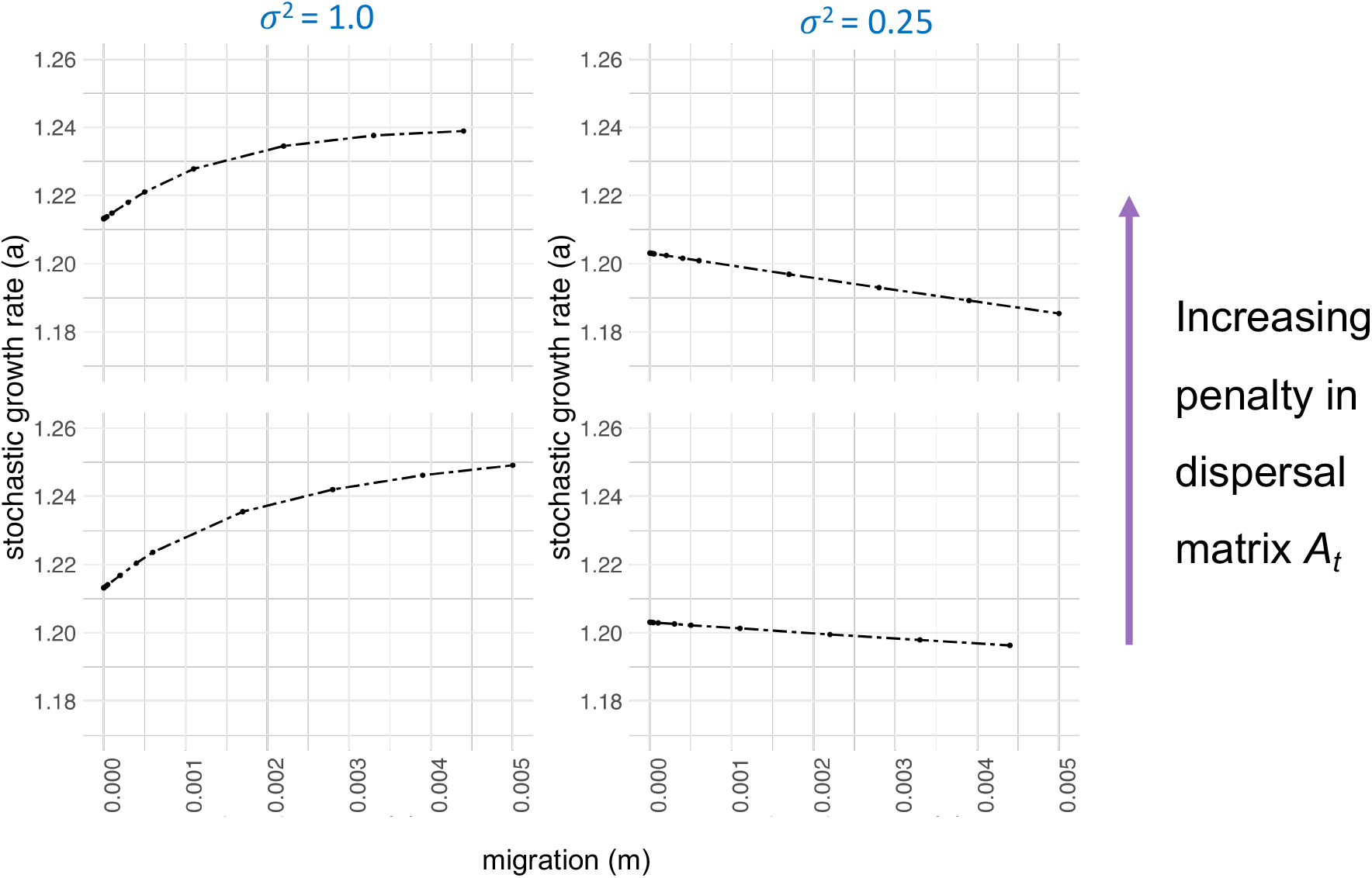
The figure has migration rate on the x-axis and the stochastic growth rate on the y-axis. The two plots in the left panel correspond to high variability (*σ*^2^ = 1) while the right panel corresponds to low variability (*σ*^2^ = 0.25). The top panel corresponds to migration penalty of 20% for a fixed 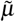 whereas the bottom panel corresponds to migration penalty of 10%. The penalty is introduced by making ∑_*i*_ **A**(*j, i*) negative by 20% and 10% as this implies individuals were lost during migration.

## 5 Main result for many sites in variable environments

We now extend the system to any number of sites. We will index the sites from 0 instead of 1. Consider sites 0, 1, …, *d* − 1 (*d >* 3) with log growth rates 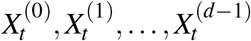 during time period *t*. The expected growth rates at each site are µ_0_, µ_1_, · · ·, µ_*d*−1_, respectively. Since we are interested in the difference in expected growth rates between of any site *j* from the unique best site 1, we define 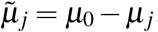 As discussed in earlier sections, the random variables 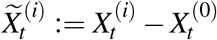 are assumed Gaussian with variance *σ*_*i*_ and de-scribe the fluctuations of the difference in growth rates at time *t*. The requirement that the log growth rates have Gaussian distribution is for simplicity of presentation; in the Appendix this assumption is relaxed to allow 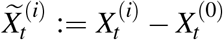 to be sub-Gaussian with variance factor τ _*j*_. The probabilistic reasoning for the theorem for multiple sites is analogous to that for the theorem for two sites.

The theorem establishes sequences of possible excursions (and their probabilities) for which the less favorable site does better than the favorable site. However, when there are more than two sites, there can be many different paths. The paths may involve no direct transition between the best site and the second-best site. Recall that the growth-rate bonus due to migration arises from small subpopulations migrating to an inferior site just as that site is experiencing an accidental run of exceptionally good conditions; and then depositing a fraction of its greatly expanded offspring back into site 0 to continue the run of normal (optimal) growth. Since migration involves only a fraction *m* of the population, the fraction making the round trip will be the even tinier *m*^2^ for a two-site system, but in general *m*^*κ*^, where *κ* is the number of migration events required to bring descendants back to site 0.

When considering a model with more than two sites we thus need to define for each site *j* a new variable *κ* _*j*_, defined as the smallest length of a cycle in migration graph ℳ that starts and ends at site 0, and passes through site *j*. For two sites, *κ* is always 2, since the possible transitions can be from site 0 to site 1 and site 1 to site 0. We incorporate *κ* by modifying the paths between sites, reflected in the dispersal matrix **A**.

### Theorem 2.

*Let* 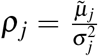 *and κ* _*j*_ *be the smallest length of a cycle in migration graph* ℳ *that starts and ends at site 0, and passes through site j. Define* <u>η</u> = min _*j*_ 2*κ* _*j*_ρ _*j*_, 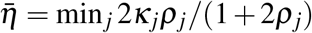. *Then for any* δ *>* 0, *we have for all m >* 0 *sufficiently small:*

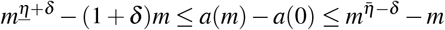

Note that the lower bound is dominated by the higher power of *m* as *m* → 0. As 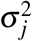 increases, the power decreases, yielding conditions that make small deviations from the case of zero migration advantageous despite the costs that are merely proportional on the order of *m*. Note that *κ* features in the two-site case as well, but since *κ* = 2 is fixed for the two site case it is absorbed as a constant.

As discussed previously, the increase of *a* near 0 is often superlinear, growing as *m*<u>^η^</u>. If the power <u>η</u> is strictly less than 1, the rapid increase in *a* near 0 will be qualitatively unaffected by a linear term for *m* sufficiently small.

### 5.1 Example of movement patterns

Consider the case for 4 sites-populations dispersing away from site 0 can return in a cycle of length 2, 3 or 4. For simplicity in the examples below we assume there is no loss of individuals. These matrices represent models where no net population is lost in migration – hence the rows sum to 0 – but nothing would be changed in the calculations below if we penalised migration. In Figure 6, the migration graphs and corresponding matrices are shown. Each graph corresponds to different length for *κ* _*j*_.

**Figure 6.**
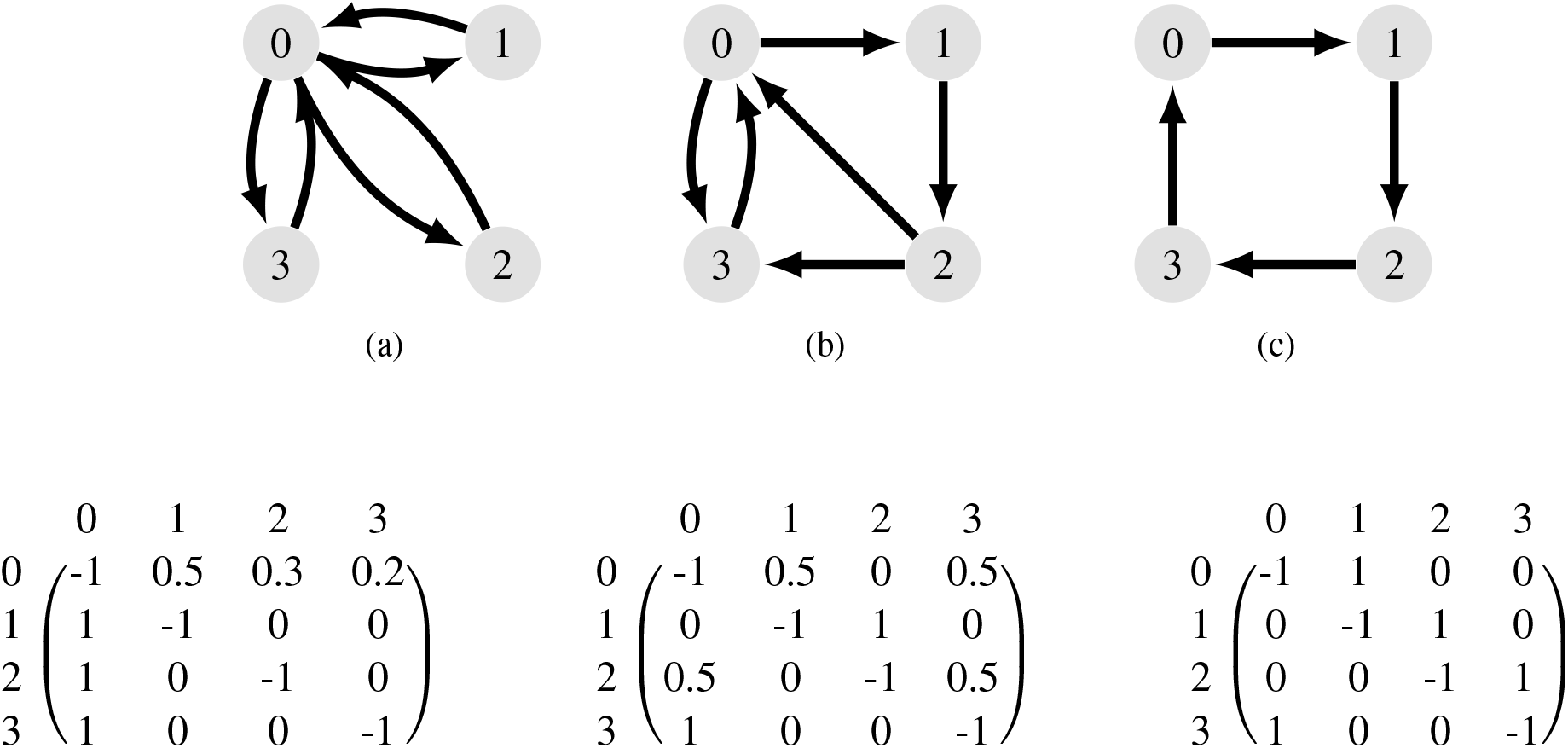
The migration graph ℳ for four-sites and their respective migration matrices. For graph (a): *κ* _*j*_ is 2 for all sites *j*, for graph (b): *κ*_1_ is 3 for site 1, and for graph (c): *κ* _*j*_ is 4 for all sites *j*.

For migration graph ℳ in Figure 7, we can calculate 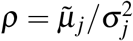 for the four-site graph. From the theorem, it is clear that to increase *a*(*m*) − *a*(0), *κ*ρ must be minimized. We see that *κ*ρ is minimized at site 1, despite the fact that it is not in the shortest cycle, nor does it have the smallest mean difference in log growth rate from the best site 0, yielding a lower bound on the sensitivity to migration on the order of *m*^0.6^. The upper bound *m*^0.41^ comes from migration to site 3, which has the lowest mean log growth rate, but compensates for this with a shorter cycle length.

**Figure 7.**
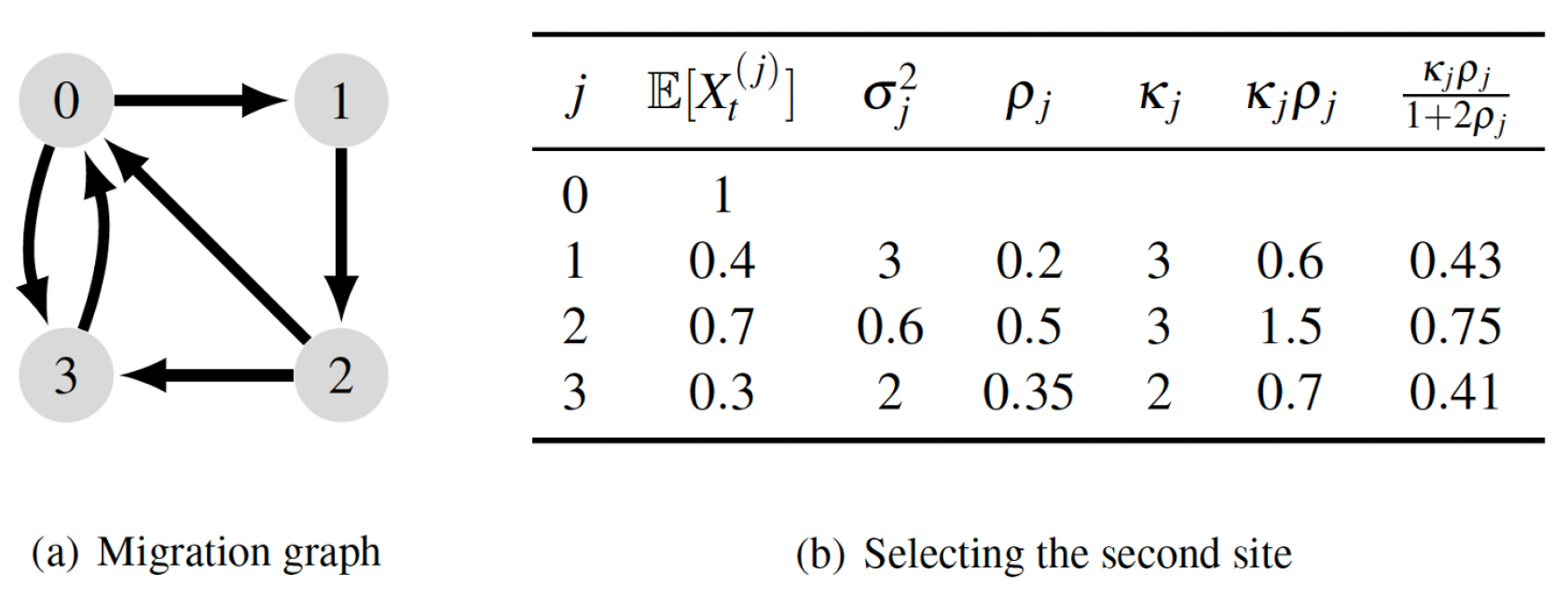
Example of calculating exponents for the migration graph given in Figure 6(b). The values of 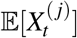 and 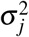 are arbitrary, and have been selected for illustrative purposes.

### 5.2 Simulations for a 4-site model

#### 5.2.1 Effect of varying fluctuations and difference in mean growth rates

For four sites (labelled 0, 1, 2, and 3), we assume site 0 has highest average growth rate µ_0_. We define the difference for sites from the best site as follows: 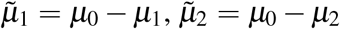 and 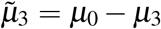 and vary the variance of the fluctuations *σ*^2^. For the simulations we consider two cases: first, we consider the vector 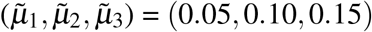. Since 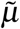 refers to differences, the vector corresponds to a case when the three sites are similar to the best site on average because the difference between sites is atmost 0.15. Now, we consider 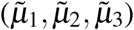 is (0.35, 0.40, 0.45) so that the difference in expected growth rate from the best site is at least 0.35. Please note that these numbers are chosen for purposes of illustration, and have no special significance. Other combinations of parameters would produce qualitatively similar results.

As shown in Figure 8, similar to the results for 2 sites, we find that increasing fluctuations increases the gain in stochastic growth rate relative to no migration. However, as the gap grows between mean growth at the best site and at the other sites, the benefits of migration for stochastic growth rate shrink, and ultimately migration will be detrimental despite high fluctuations. Hence populations cannot benefit from variability in this case (last panel Figure 8). We consistently find that the smallest kappa yields the greatest increase in stochastic growth rate, as discussed in the next section.

**Figure 8.**
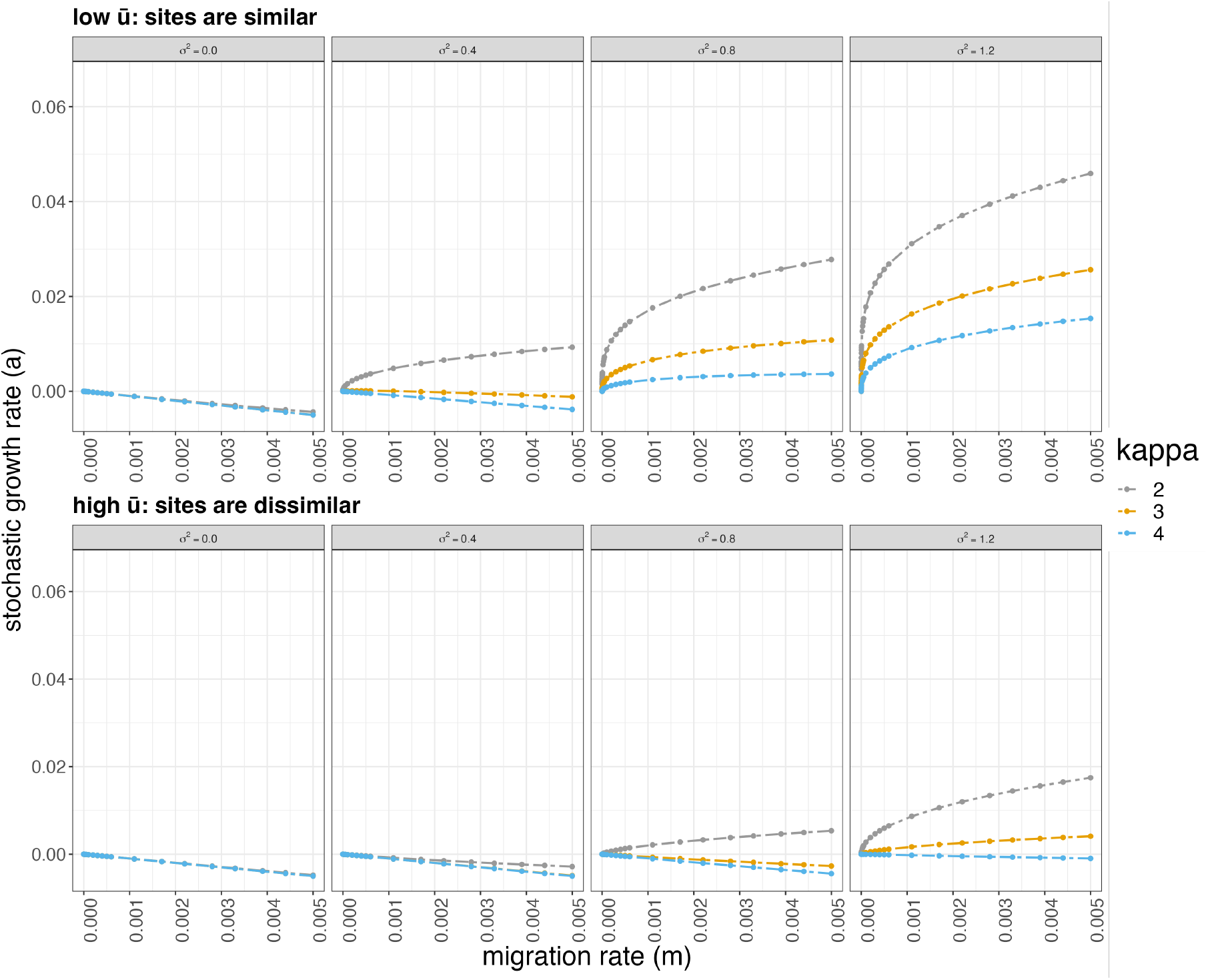
The x-axis is migration rate and y-axis is the stochastic growth rate: *a*(*m*). The top panel corresponds to values for the vector 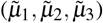 such that the sites are similar. The panel on the bottom corresponds to values such that sites are dissimilar. Each color corresponds to a value of *κ* (for four sites) since an excursion can be made in steps of 2, 3 and 4.

#### 5.2.2 Effect of varying path length between sites

We now examine how *κ* affects the long-term growth rate. Recall that *κ* _*j*_ is defined to be the length of shortest cycle in the migration graph that starts and ends at 0, passing through *j*.

In Figure 8, each color corresponds to a value of *κ* for four sites since an excursion can be made in lengths of 2, 3 and 4. From the theorem, we expected the shortest excursion with *κ* = 2 to lead to the maximum gain in stochastic growth rate, and that is what we find in Figure 8. The longest excursion with *κ* = 4 leads to the lowest increase in stochastic growth rate. Although populations sample more variability by increasing the number of possible paths and sites, it is important to return to the best site in the shortest sequence possible to maximize the gain in overall growth rate. That is, increase in growth rate is determined by being able to gain access to the best site and return with a smaller number of low-probability migration events (lower *κ*).

In addition, as evidenced by the last two panels in Figure 8, the length of *κ* matters when the fluctuations are high and when the sites are more similar to the best site. The bottom two panels of Figure 9 show how the stochastic growth rates for different path lengths remain similar when the difference between sites is increased (high µ). Hence, path length matters most when the sites are similar, as illustrated in top two panels of Figure 9.

**Figure 9.**
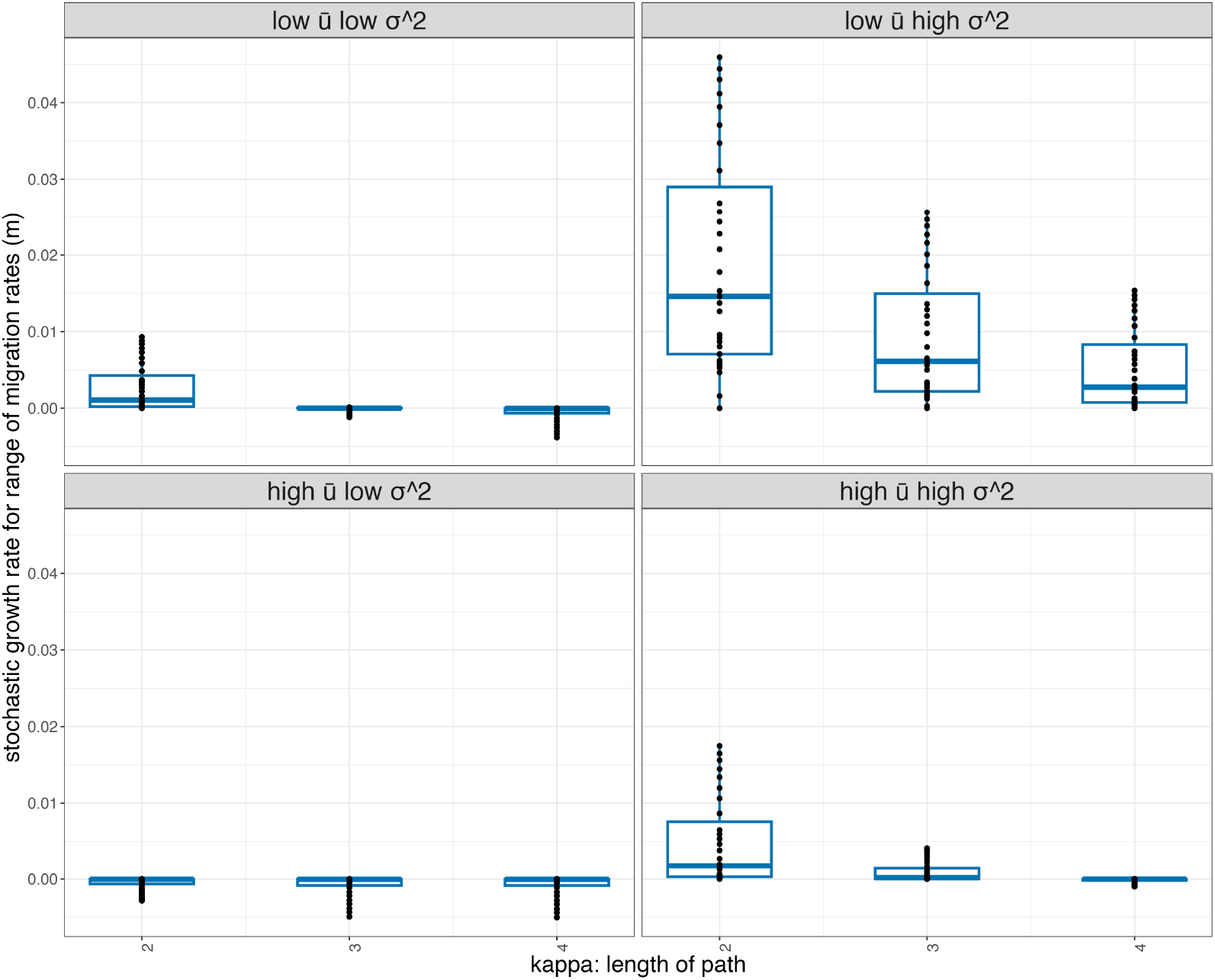
The effect of changing topology on growth rates in a 4-site model. The x-axis is the length of the shortest path to the best site (site 0) in steps of 2, 3 and 4. The y-axis are the values of stochastic growth rate for a range of *m*. Each panel corresponds to a fixed level of *σ*^2^, and varies in low and high difference in expected growth rates from best site.

### 6 Discussion

We provide conditions under which introducing a small amount of migration will increase the growth rate of the population relative to the zero-migration case. Our research contributes fundamentally to the literature by proving that the increase in long-run growth rate *a*(*m*) − *a*(0) with migration at rate *m* is bounded above and below by powers of *m* for *m* in a neighborhood of 0, in a simple model of multi-site population growth that takes environmental variability into account. These powers depend on the ratio 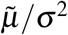, where 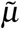 is the difference in expected growth rates and *σ*^2^ is the variance of the difference of site-specific growth rates at time *t*. Our results complement the analysis in [20] of optimal migration rates for populations with stochastic growth rates at different sites.

We illustrate our results (through simulations) by examining a wide range of fluctuations, differences in average growth rates between sites, and path lengths. We find that as the average (over time) qualities of the sites diverge, the increase in long-run growth rate due to introduction of migration becomes less pronounced. Environmental fluctuations compensate (to an extent) for the fixed difference in mean log growth rates. Biologically, this means that habitat quality and fluctuations are both important determinants of migration evolution and population persistence.

By changing how each site is connected to the ‘best’ site (varying *κ*), we find that when variability *σ*^2^ is low, excursions of any length are unlikely to overcome the deficit in average growth (refer to Figure 9). When variability *σ*^2^ is high, excursions may overcome average differences when the length of *κ* is small and will yield the maximum increase in stochastic growth rate in that case. Recent studies on seabirds [19, 33] show that a longer migration route does not necessarily have carry-over effects on subsequent reproduction and apparent survival for little auk *(Alle alle)* and lesser black-backed gulls *(Larus fuscus)*, respectively.

Our results build on earlier work that emphasized that the role of migration in allowing populations to keep pace with spatial and temporal variation in the habitat [53, 8, 61]. [5] formulated the concept of “spreading of risk” as a way to survive and reproduce in a spatially variable environment so that any one failure will not be decisively harmful. This concept recognizes the importance of migration as an escape from unfavorable conditions, and adaptation to changing environments by exploiting new sites. [53] intuited that the rate of migration should be highest in species inhabiting the most temporary habitats and found that migratory movement in terrestrial Arthropoda is positively correlated with impermanence of the habitat.

Numerous empirical studies have found that dispersal is more common in rapidly-changing habitats; e.g., [18] studied 35 species of plant hoppers living in habitats of varying persistence, and found that winged morphs were more common in ephemeral habitats while flightlessness was favored in persistent habitats.

A similar argument holds in the case of diapause, which is a delay in biological development, often at an early life stage. For example, many plant species produce seeds that do not germinate immediately, but remain dormant for extended times forming seed banks [60, 61]. The paper by [54] considers a two-year life-cycle model of a species, where the first year is spent growing from a seed to an adult, and the second year is spent producing seeds. They examined the effect on growth rates if a small fraction of the seeds remains dormant for one year. When vital rates remain constant over time, delaying individual growth reduces population growth. However, when environmental conditions are variable, even a tiny amount of dormancy can boost population growth. This finding parallels ours — it is equivalent to the present model with all sites having exactly the same average growth rate — but requires a substantially different analysis. Taken together, these results determine conditions under which there are departures from the reduction principle, that is, the selection for reduced dispersal (under constant selection) [3, 4].

Our findings have implications in this era of global habitat degradation, altitudinal range-shifts, and species redistribution [37, 25] which provide directions for future research. Importantly, our work provides a way to study migration patterns (of birds or mammals) by examining variation in environmental conditions, and metrics describing site-specific growth rates. These results may also be important in conservation biology and low-density populations: when there are superior sites in a sea of poor habitats, variability, habitat quality across space, and length of path may be key to determining the importance of migration.

Theoretical work on the evolution of dispersal has a long history [27, 39, 43, 28, 56, 2]. Indeed, there is a large and rich literature in dispersal and metapopulation models [56, 12, 13, 55, 36, 26, 45, 47] that remain important in most applications. Studies that unify theory and experiment, as discussed by [40], will provide ways to test or parameterize theoretical models [49]. It is pertinent to account for density-dependence when there are variations in population growth rates across space and analyze how adding more individuals that migrate impacts stochastic growth rate. Importantly, our research does not distinguish between different patterns of movement, such for breeding dispersal or seasonal migratory patterns and will be an important avenue for further research [41, 11]. Finally, future work may also examine how variability in dispersal rate impacts overall populations, and account for structure in spatial sites.

## 7 Conclusions

We solve a fundamental question in evolutionary ecology: can dispersal between sites evolve in variable environments? We use a simple model in which one site is on average the best (highest expected growth rate), and dispersal moves individuals between the best site and other sites. We show that sufficient variability means that an individual who disperses can exploit favorable runs of the environment in other sites. We provide precise inequalities that demonstrate the trade-off between differences in expected growth rates between sites, and variability between sites. We illustrate our results (with simulations) for two and four sites. For the multiple (≥ 2) site case, we consider the effect of alternative paths to return to the best site.

This fundamental idea about the evolution of dispersal is old but had not been demonstrated mathematically even in a simple model.

## Contributions

HJ assisted with the mathematics, planned and carried out the simulations, and was primarily responsible for writing the paper. DS conceived and carried out the mathematics, and assisted with the simulations. ST proposed, led, and assisted with all phases of the project. All authors contributed to shaping the final version.

## Acknowledgement

HJ would like to thank Siddharth Srinivasan for the very helpful discussions that enabled a better understanding of the mathematics and large deviation theory. HJ would also like to thank Nikunj Goel for providing constructive feedback for editing the draft. We thank Stanford University and University of Oxford as our funding sources.

## Appendix A

### Approximately linear relationship near 0 in the Log-Log Plot for stochastic growth rate (a) versus migration rate (m)

**Figure S1.**
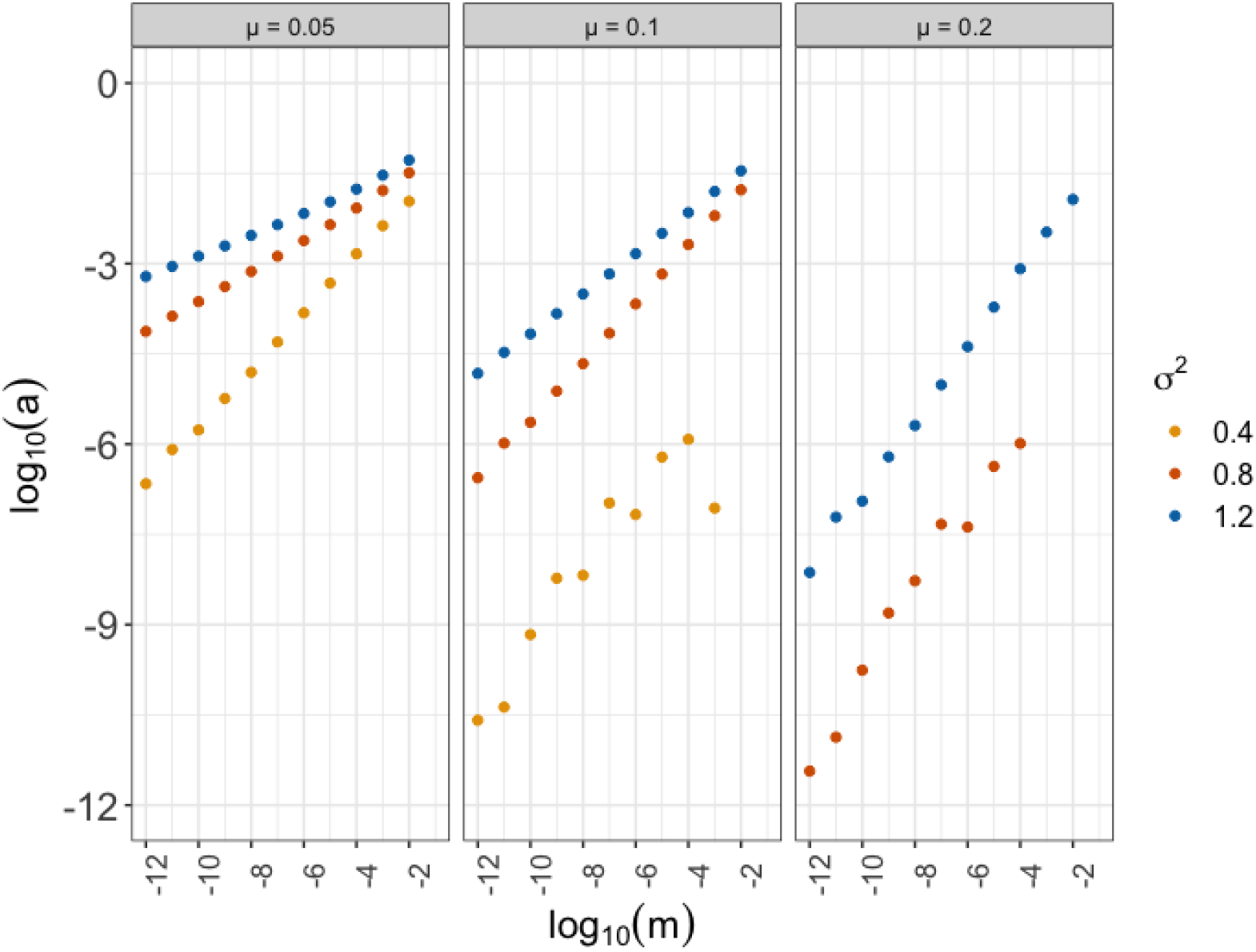
The y-axis represents logarithm of the stochastic growth rate (*a*). On the x-axis is the logarithm of the migration rate (*m*). Each veritcal panel corresponds to expected growth rates between sites 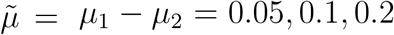 Colours indicate values of variability, *σ*^2^. There is no value for *σ*^2^ = 0 as the stochastic growth rate (*a*) is negative and thus the logarithm for it is not defined.

### Simulated slopes within bounds from the Theorem

**Figure S2.**
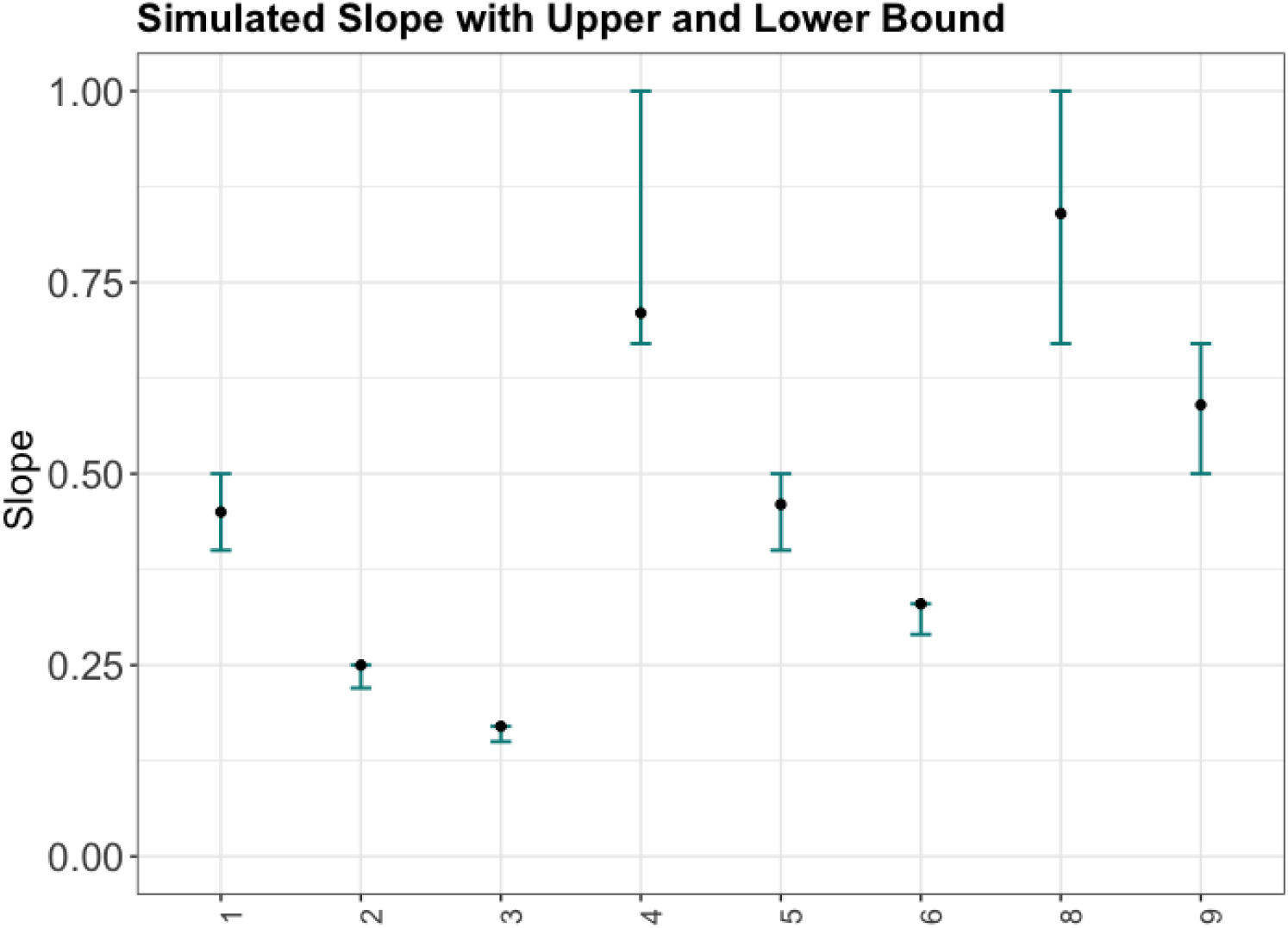
The y-axis represents the slope, upper bound and lower bound for different combinations of *µ* and *σ*^2^ shown in Figure 2. The slope (equivalent to the power of *m*) is calculated between two values of migration rate (*m*_1_ = 1 × 10^*−*12^ and *m*_2_ = 1 × 10^*−*10^) as 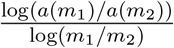. The blue-green bars represent the upper and lower bounds predicted by the theorem asymptotically as log *m* → −∞, and the black points represent the simulated slope value. The x-axis refers to different combinations of 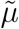 and *σ*^2^ as defined in Appendix Figure 1, and tabulated in Table 1. Note that for simulation 7 the power is greater than 1, hence the change in growth rate is negative, hence there is no log plot and no slope.

**Table 1.** Table for slopes and corresponding upper and lower bounds.

| Sim No. | $\tilde{\mu}$ | $\sigma^2$ | Slope | Upper bound | Lower bound |
| --- | --- | --- | --- | --- | --- |
| 1 | 0.05 | 0.40 | 0.448 | 0.500 | 0.400 |
| 2 | 0.05 | 0.80 | 0.247 | 0.250 | 0.222 |
| 3 | 0.05 | 1.20 | 0.169 | 0.167 | 0.154 |
| 4 | 0.10 | 0.40 | 0.711 | 1.000 | 0.167 |
| 5 | 0.10 | 0.80 | 0.460 | 0.500 | 0.400 |
| 6 | 0.10 | 1.20 | 0.328 | 0.333 | 0.286 |
| 7 | 0.20 | 0.40 | NA | 2.000 | 1.000 |
| 8 | 0.20 | 0.80 | 0.838 | 1.000 | 0.667 |
| 9 | 0.20 | 1.20 | 0.592 | 0.667 | 0.500 |

## Appendix B

The goal of the appendix is to describe and prove Theorem 1 and 2 discussed in the main manuscript. Note that for simplicity, the theorems for the case of two sites and more than two sites are presented separately in the main manuscript. In the Appendix, we present the general theorem for any number of sites and the 2-site theorem is a special case of the general theorem. First, we state the key notations and assumptions. We define sub-Gaussian random variables and state Theorem 1 and 2.1. Next we define *ancestries* and *excursions* — sequences of sites that are allowed by the migration graph — that will be enumerated in the derivation of upper (in section 4) and lower bounds (in section 5).

We summarize some key assumptions and definitions from the main manuscript below:

### 1. Notation and basic assumptions

- Let there be *d* sites such that populations 0, …, *d* − 1 have log growth rates 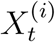 during time period *t* at site *i*. The vectors 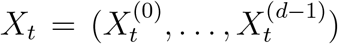 are assumed independent, but the components within each *X*_*t*_ — corresponding to growth rates at different sites at the same time — may be dependent.
- We write 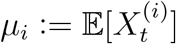, and assume that *µ*_0_ is strictly the largest of these.Writing 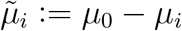, these are all positive. We write **X** for the complete collection of all 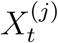 for *j* = 0, 1, …, *d* − 1, 0 ≤ *t <* ∞.
- The random variables 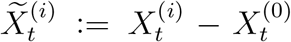 are assumed sub-Gaussian with variance factor *τ*_*i*_, defined below in (2). When they are Gaussian, as we have assumed for the simulations, *τ*_*i*_ is then simply the variance.
- Suppose *G*_1_, *G*_2_, … is an i.i.d. sequence of *d*×*d* diagonal matrices. The diagonal elements of *G*_*t*_ are the log growth rates at each site at time *t* such that 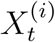 all have finite mean *µ*_*i*_ and finite variance.
- We let *A*_*t*_ be an i.i.d. sequence of *d* × *d* matrices with negative entries on the diagonal and non-negative entries off the diagonal. We assume that the pairs (*G*_*t*_, *A*_*t*_) are all independent, but not necessarily that *A*_*t*_ and *G*_*t*_ are independent for any given *t*. The entries of *A*_*t*_ are all assumed to be almost-surely bounded.

#### 1.1. Migration graph

We define the migration graph ℳ to be an irreducible directed graph whose vertices are the sites {0, …, *d* − 1}, representing the transitions that have nonzero probability. We assume that ℳ is connected.

Let *m* parameterize the overall rate of migration. Thus, migration rates from site *i* to site *j* in *mA*_*t*_(*j, i*). Population distributions are thus column vectors, and are updated from time *t* − 1 to time *t* by left multiplication.

When *i* → *j* (*i* is directly linked to *j* in M) we assume 𝔼[(log *A*_*t*_(*j, i*))_+_] exists and is finite, though *A*_*t*_(*j, i*) may still have nonzero probability of being 0. We write *µ*_*A*_ := max_*i,j*_ 𝔼[(log *A*_*t*_(*j, i*))_+_].

We will assume that *A*_*t*_(*i, i*) is almost surely bounded below, and we will henceforth always be assuming that *m* is sufficiently small that 1 + *mA*_*t*_(*i, i*) *>* 0 for all *i* with probability 1. We write Δ^*i*^ := −𝔼[*A*_*t*_(*i, i*)], which we assume also to be finite.

Using the system of equations defined in the main manuscript in Equation (3), we define

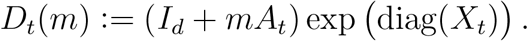

For *m >* 0 the i.i.d. sequence *D*_*t*_(*m*) satisfies the conditions for the existence of a stochastic growth rate independent of starting condition [Coh79]. That is, if we define the partial products

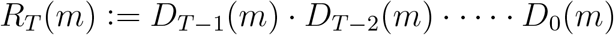

then

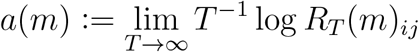

are well defined deterministic quantities, in the sense that the limit exists almost surely, is almost-surely constant, and is the same for any 0 ≤ *i, j* ≤ *d* − 1.

The quantity we are trying to approximate and provide the bounds for is

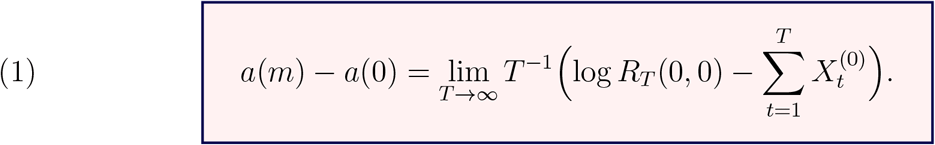

#### 1.2. Sub-Gaussian variables

Although we describe our results using Gaussian random variables, the results are more general and apply to sub-Gaussian random variables. The random variables 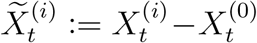 are assumed sub-Gaussian with variance factor *τ*_*i*_. This is a bigger class of distributions which includes Gaussian distribution. Please note that in the main document, all simulations are done assuming Gaussian.

A random variable *Z* is said to be *sub-Gaussian* if it has finite *variance factor* (the term applied in [BLM13]) *τ* = *τ* (*Z*), defined by

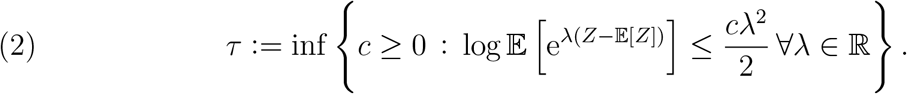

(*τ* (*Z*) is called the *sub-Gaussian standard* in [BK00].)

### Biological Intuition

This may be thought of as an upper bound on the scale of the tails, and it is this that determines the upper bound on the sensitivity of the stochastic growth rate to migration.

Note that the Legendre transform for the moment generating function *I*_*Z*_(*z*) := sup_*λ*_{*λz*− log E[e^*λ*(*Z−*𝔼[*Z*])^]} satisfies *I*_*Z*_(*z*) ≥ *z*^2^*/*2*τ* (*Z*) for all *z*. Applying Markov’s inequality implies the standard sub-Gaussian concentration inequality

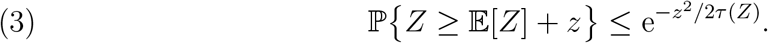

To describe the lower bounds we define a lower subvariance *<u>τ</u>* (*Z*) by

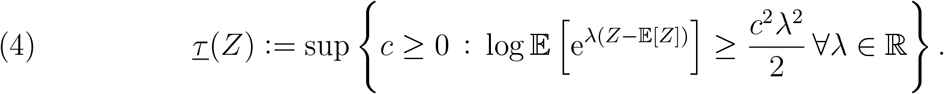

Then for all *z* we have *I*_*Z*_(*z*) ≤ *z*^2^*/*2*τ* (*Z*). We define also

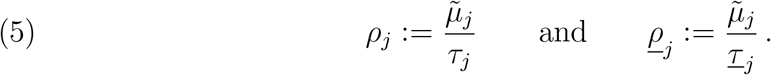

If *Z* is Gaussian with variance *σ*^2^ then of course *τ* (*Z*) = *<u>τ</u>* (*Z*) = *σ*. The calculation of *ρ* is illustrated in Figure 3.

**Figure S3.**
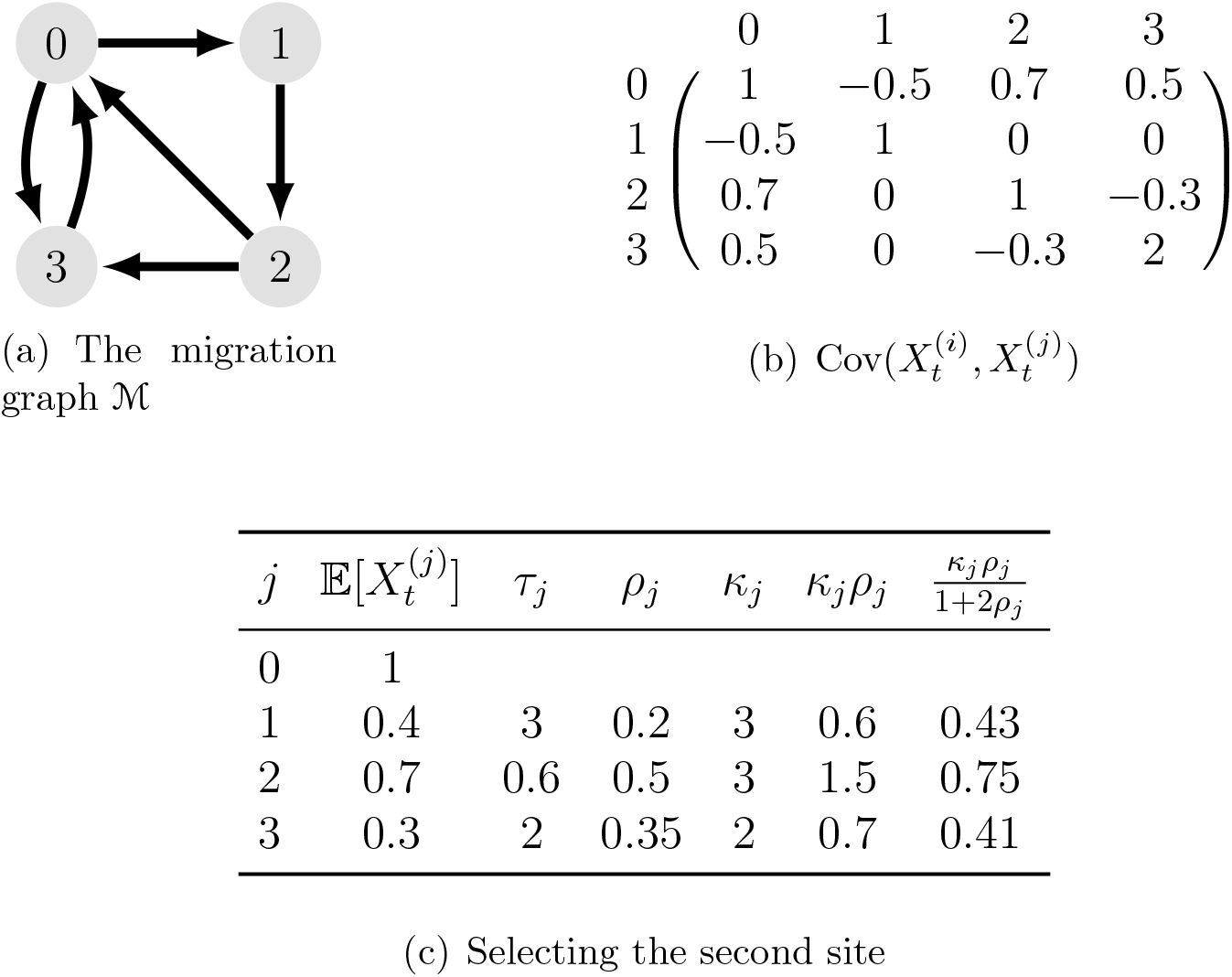
Calculating *ρ* on a four-site graph. For simplicity, we assume the variables *X*_*t*_ are all Gaussian, so that *τ*^2^ = *τ*^2^ is the variance, and *ρ* = *ρ*. We see that *κρ* is minimized at site 1, despite the fact that it is not in the shortest cycle, nor does it have the smallest mean difference in log growth rate from the optimal site 0, yielding a lower bound on the sensitivity to migration on the order of *m*^0.6^. The upper bound *m*^0.41^ comes from migration to site 3, which has lower mean log growth rate, but compensates for this with a shorter cycle length.

Like variances, sub-Gaussian variance factors are sub-additive for sums of independent random variables, as stated in [BK00, Lemma 1.7]: If *Z*_1_, …, *Z*_*k*_ are independent random sub-Gaussian random variables with variance factors *τ*_1_, …, *τ*_*k*_. Then *Z*_1_ +· · · +*Z*_*k*_ is sub-Gaussian with variance factor less than or equal to *τ*_1_ + · · · + *τ*_*k*_.

### 2. Theorem

In the Appendix, we present the general theorem for any number of sites and the two-site as a special case after the describing the theorem. For simplicity, the theorems for the case of two sites and more than two sites are presented separately in the main manuscript. This is because when there are more than two sites (*d >* 2), the result is complicated for two reasons: First, the effect of migration will be dominated by the site that has the second fastest growth; second, the increment to growth will be smaller if direct transition between the two leading sites is impossible. For this purpose, for each 1 ≤ *j* ≤ *d* − 1 we define *κ*_*j*_ to be the smallest length of a cycle in M that starts and ends at 0, and passes through *j*. (Thus 2*d* − 2 ≥ *κ*_*j*_ ≥ 2, and *κ*_*j*_ is equal to 2 when there are direct paths from 0 to *j* and from *j* to 0.)

#### Theorem 1.

*Under the assumptions of section 1, let* 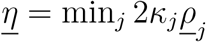, *and* 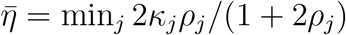 *Then for any δ >* 0:

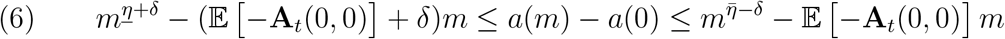

*for m >* 0 *sufficiently small*.

Note that the upper bound is dominated by the lower power of *m*, and the lower bound is dominated by the higher power of *m* and *κ*_*j*_ is the smallest length of a cycle in M that starts and ends at 0, and passes through *j*. From the theorem, it is clear that to increase 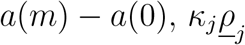 must be minimized.

Since the dispersal matrix is not constant the linear term has a coefficient that is the expectation of **A**_*t*_(0, 0). In the main manuscript, this is just −1. (It has been written as −𝔼 [−**A**_*t*_(0, 0)] to make clear that it is generally understood to be negative.

The important point is that the effect of the diagonal terms in **A**_*t*_ is always nearly linear in *m*, while the increase of *a* near 0 is often superlinear, growing as a power of *m* smaller than 1. If the power 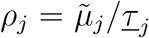 is strictly less than 1, the rapid increase in *a* near 0 will be qualitatively unaffected by a linear term for *m* sufficiently small. Even when the linear term is negative (as we will generally be assuming it to be), the growth rate *a* will still be increasing on a small interval of *m >* 0.

The cost of migration is governed by the row sums of matrix **A**_*t*_. Since individuals who migrate are lost from the originating site, it would be reasonable to require that the total of all migration from a site ∑_*i*_ *A*_*t*_(*j, i*) is no more than 0; if individuals are lost in the process of migration then the sum will be negative. For the section on migration penalty in the main manuscript, we make the ∑*i A*_*t*_(*j, i*) negative to account for loss of individuals.

#### 2.1. Case when there are 2 sites

This is a special case for the Theorem above since *κ* = 2 when there are only two sites. In this case, the difference between stochastic growth rates, *a*(*m*) and *a*(0) is bounded below and above (scaled by a constant) as follows: For any positive *δ*,

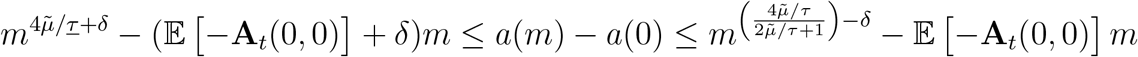

for positive *m* sufficiently close to 0.

Now that the theorems have been stated, we will define the excursion decomposition that will be used to prove the bounds for *a*(*m*) − *a*(0).

### 3. Excursion decompositions (path length)

Since we are assuming the unique maximum average growth rate is at site 0, the maximum growth for the perturbed process will arise from rare excursions away from 0; in particular, from those that include the (not necessarily unique) site that minimises *ρ*κ** in (5).

Define E to be the set — called the *excursions* — of cycles in the migration graph that start and end at 0, with no intervening returns to 0. For an excursion **e** we write |**e**| for the length of the cycle minus 2 — that is, the number of time steps spent away from 0.

As is standard in statistical mechanics, the change in stochastic growth rate will ultimately be determined by the tradeoff between the penalty for migration and for spending time away from the optimal site, and the entropy gain from increasing the number of possible paths as the number of allowed transitions grows. To calculate this tradeoff quantitatively we need a notation that will allow us to enumerate the paths — defined by a sequence of excursions — that will draw similar penalties. Such a sequence may be thought of as an *ancestry*, as it records the locations of the ancestors of a generation *T* individual in each of the preceding generations. For purposes of enumeration the essential features will be the number of excursions in the ancestry, total length of the excursions, and the number of transitions between sites.

For a given excursion **e**, comprising sites 0 = **e**_0_, **e**_1_, …, **e**_|**e**|*−*1_, **e**_|**e**|_, **e**_|**e**|+1_ = 0, we define

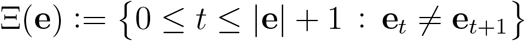

#### Biological Interpretation

Ξ(**e**) denotes the set of change points in an excursion, i.e, it collects the times at which a transition is made, meaning *e*_*t*_≠ *e*_*t*+1_. The total number of transitions made from one site to the next, i.e, #Ξ(**e**) is denoted by *s*.

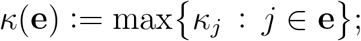

#### Biological Interpretation

Here *κ*_*j*_=min {no. of edges of *P* |*P* is a closed path passing through 0 and *j*}. That is, *κ*_*j*_ is the length of the smallest cycle that passes through vertex *j* and starts and ends at 0. The smallest number of steps possible in a cycle containing *j* and 0 can not be strictly greater than the actual number of transitions in an excursion passing through *j*, which defines such a cycle. Therefore, *κ*(**e**) ≤ #Ξ(**e**) = *s* since *κ*(**e**) is the maximum of all *κ*_*j*_ for *j* ∈ **e**. We call *κ*(**e**) the diameter of *ê*.

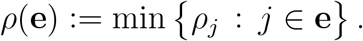

#### Biological Interpretation

This is the loss in expected log growth at the site that might be thought of as the “target” of the excursion — the site where the loss is the smallest, scaled by the variance, which turns out to be the appropriate combination for measuring the influence of the fluctuations at the non-optimal site *j*, given that an excursion reaches that site.

We write 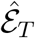 for the collection of ancestries — sequences of excursions — that can be fit into time {1, …, *T*}. That is, an element 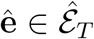 has an *excursion count k*(**ê**), such that each *i* ∈ {1, …, *k*(**ê**)} there is a pair (*t*_*i*_, **ê**_*i*_) with *t*_*i*_ ∈ {2, …, *T* − 1} and 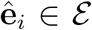 satisfying

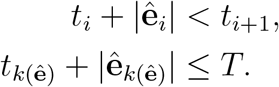

*t*_*i*_ may be understood as the starting time of excursion *i*, which then continues for |**ê**_*i*_|, and so is definitely back at 0 before the next excursion starts at time *t*_*i*+1_. We write the total length of an ancestry as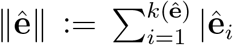. We also write 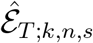 ^for^ the subset of 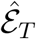 comprising ancestries whose excursion count is *k*, whose total length is *n*, and the sum of whose change-point counts #Ξ(**ê**_*i*_) is *s*. That is, *s* = #Ξ(**ê**), where 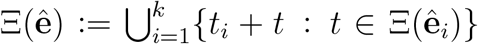. The *null ancestry* is the element of 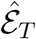 with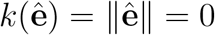.

The (0, 0) entry of the product *R*_*T*_ will be a sum of terms that are enumerated by elements of 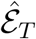, corresponding to paths through the sites. We define new random variables as a function of the realizations of **X** and of **A** (the collection of all matrices *A*)

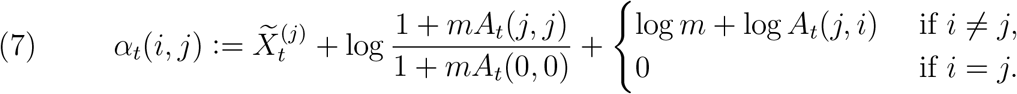

Given an excursion **e** and a starting time *t*_0_ ∈ {0, …, *T* − |**e**| − 1} we define the rando variables

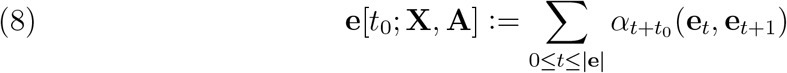

Of course, this sum may be −∞, if it includes a transition at which the corresponding entry of *A* is 0. But our assumptions imply that it is finite with nonzero probability if **e** ∈ E. Given an ancestry 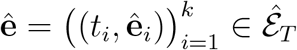 we define

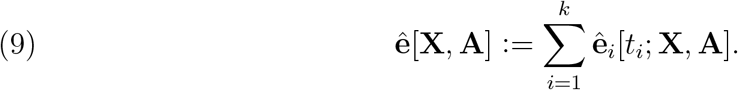

To find the bounds for equation (1), we will use Lemma 2 to find an upper bound for log *R*_*T*_ (0, 0). The proof invokes summing over ancestries.

**Figure S4.**
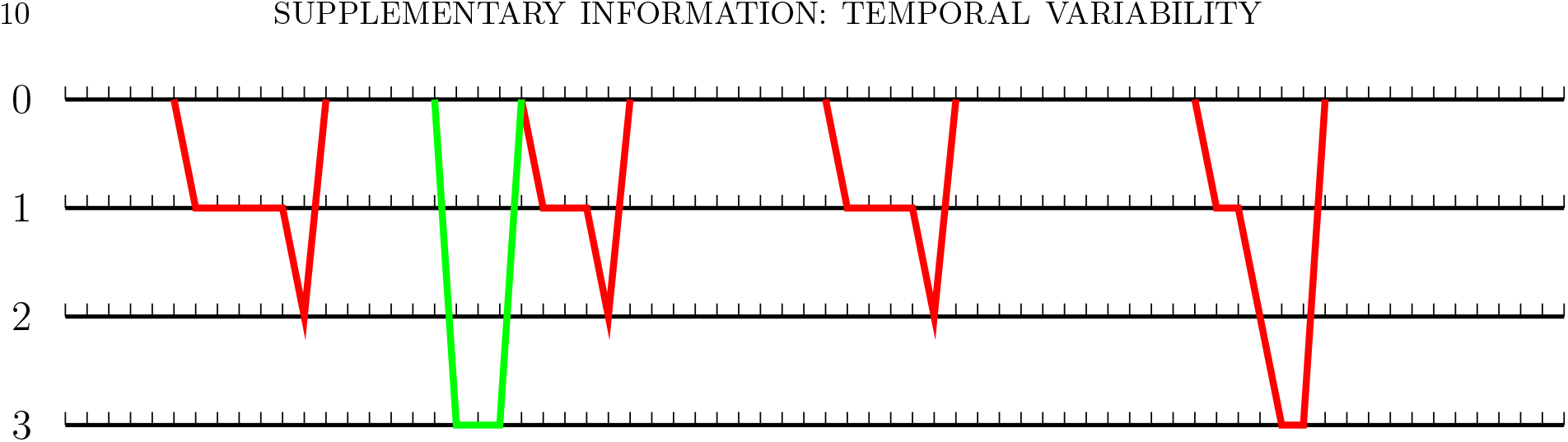
An ancestry for *T* = 70 comprising *k* = 5 excursions. This is based on the migration graph example from Figure 3. Three excursions (red) have diameter 3, and one (green) has diameter 2. Note that one timepoint (*t* = 21) is included in two different excursions. The red excursions all have *ρ*(**ê**) = 0.2, and the green excursion has *ρ*(**ê**) = 0.35. The lengths are 6, 3, 4, 5, 5, giving the sequence a total length *n* = 23. The change-point counts are 3, 2, 3, 3, 4, summing to *s* = 15.

##### Lemma 2

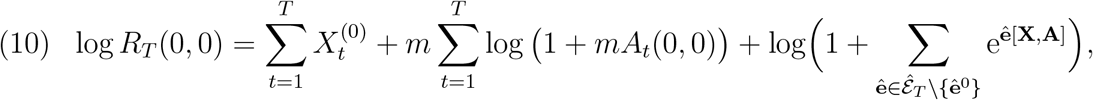

*where* **ê**^0^ *is the null ancestry*.

*Proof*. We have, by definition,

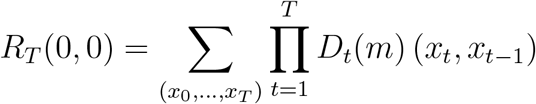

where the summation is over (*x*_0_, …, *x*_*T*_) ∈ {0, …, *d* − 1}^*T* +1^ with *x*_0_ = *x*_*T*_ = 0. We may restrict the summation to (*T* +1)-tuples such that *D*_*t*_(*m*) (*x*_*t*_, *x*_*t−*1_) *>* 0, which will only be true when (*x*_*t−*1_, *x*_*t*_) is an edge of ℳ. Such sequences of states map one-to-one onto ancestries.

By the definition of matrix products,

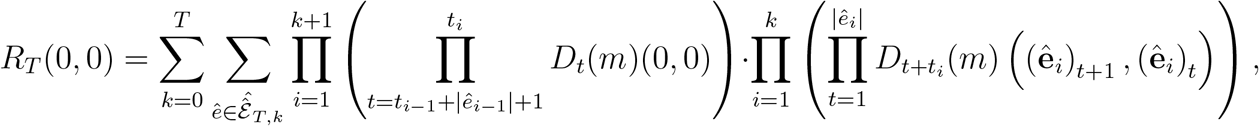

where *t*_0_ = 0 and *t*_*k+1*_ = *T*. The logarithm of the summand in the above expression is

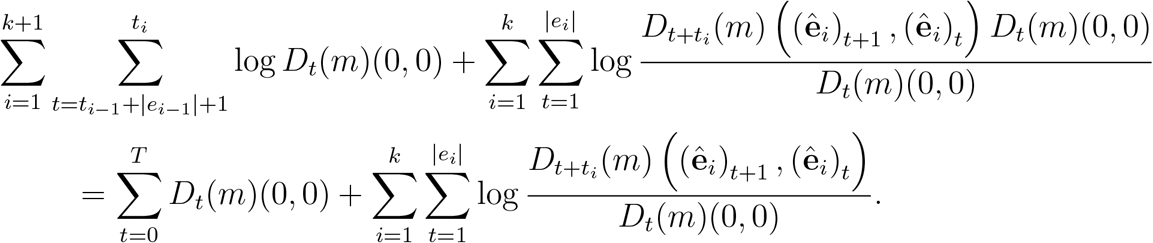

We have 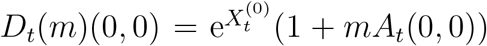. Taking the logarithm of the expression, we get for the expression above

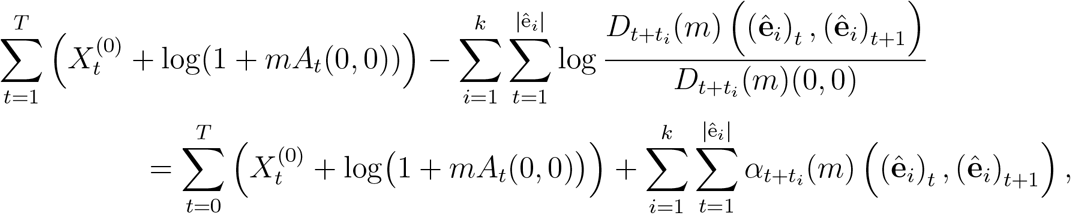

since

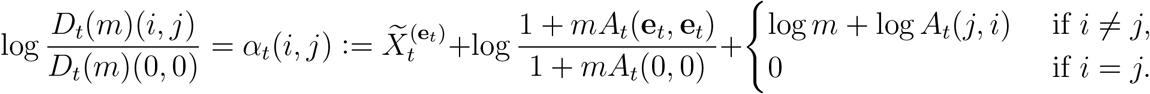

The result then follows from splitting off the null excursion and invoking the definition of the random variable **ê**[*X, A*]. □

We have then

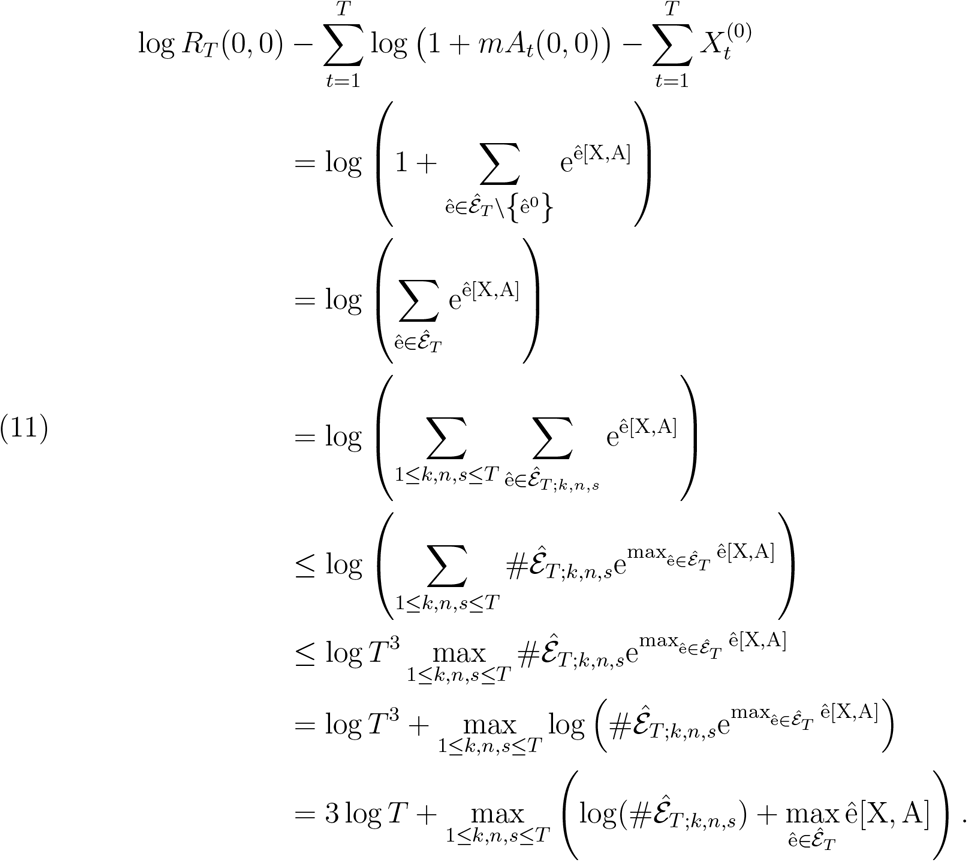

By the Strong Law of Large Numbers,

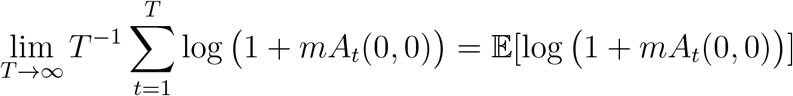 almost surely.

As we have assumed *A*_*t*_(0, 0) to be almost-surely bounded, we have for any positive *δ*

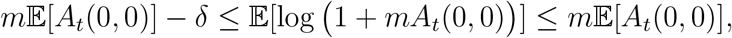

for *m* sufficiently small.

Since

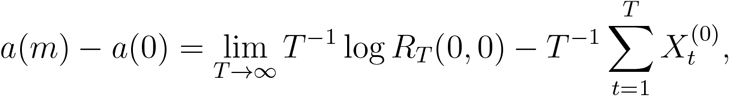

we have the bounds

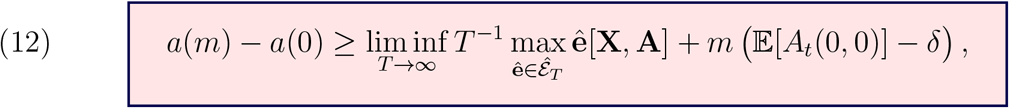

and

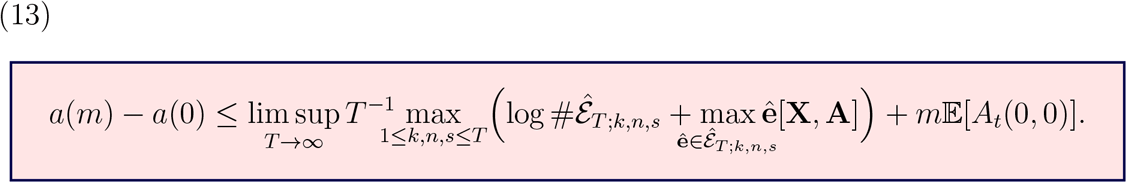

### 4. Derivation of the upper bound

We prove the upper bound in (6). We may replace *A*_*t*_(*i, j*) by *A*_*t*_(*i, j*) ∨ 1 for any (*j, i*) ∈ ℳ, since decreasing *A*_*t*_ can only decrease *a*(*m*) − *a*(0). That is, we put a floor under those off-diagonal elements which are allowable migrations. This avoids the nuisance of having entries be sometimes 0, and an upper bound that holds under these conditions will hold *a* fortiori under the original conditions. Indeed, we may assume without loss of generality that all *A*_*t*_(*j, i*) = 1 identically for *i*≠ *j*, since *a*(*m*) — the stochastic growth rate with the true values of *A*_*t*_ — is no larger than *a*(*A, m*; 1), the stochastic growth rate where all values of *A*_*t*_ that are permitted to be nonzero are replaced by 1. This changes our upper bound only by a constant, which may be absorbed into the constant of the theorem. Thus, we will proceed under this assumption.

An element of 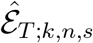 may be determined by the following choices:

i. Choose *k* points out of *T* where the excursions begin, yielding no more than 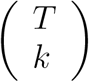 possibilities;
ii. Choose *k* numbers for the lengths of the excursions that add up to *n*, yielding*k* no more than 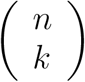 possibilities;
iii. Choose *s* − 2*k* time points within these excursions as times when there is a change of site, yielding at most 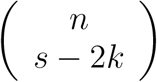 possibilities;
iv. There are no more than *d*^*s*^ ways to choose the sites to which the excursions move at the *s* times when there is a change.

Then using the points above with the a crude bound based on Stirling’s Formula (derived in section 6.1), we get:

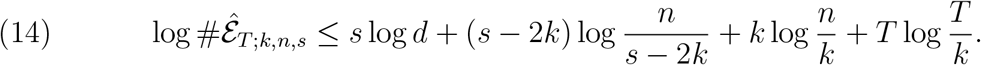

#### Claim 3.

*Suppose that ρ*_*j*_ *and *κ**_*j*_ *are each minimised at site j* = 1. *For any δ >* 0 *and*

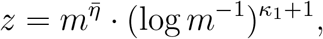

*we have*

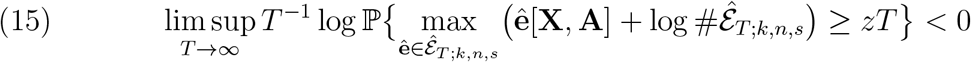

*for m >* 0 *sufficiently small, meaning m* ∈ (0, *m*_0_) *where m*_0_ *is a constant, depending only on the model parameters and δ*.

The proof of Claim 3 is provided at the end of the section for Derivation of the Lower Bound 5. Assuming the Claim,

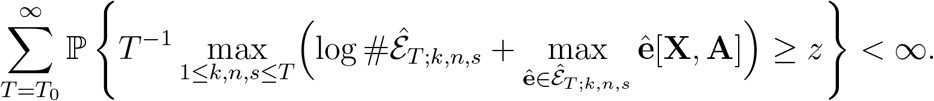

By the Borel–Cantelli Lemma, this implies that with probability 1 this event occurs only finitely often. It follows that the limsup is no greater than *z* almost surely, and hence, by (13), that

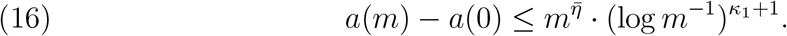

It remains only to clear away the assumption that that *κ*_*j*_ and *ρ*_*j*_ are both minimized at site 1. We do this by stratifying the excursions further by their diameter (recall the definition from section 3). Define

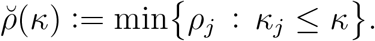

If **e** is an excursion with diameter *κ*, then any site *j* included in **e** has *κ*_*j*_ ≤ *κ*, hence also 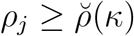. Furthermore,

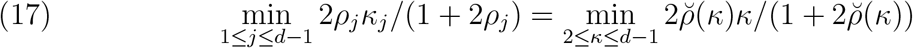

The maximum in (13) may be written as a maximum over (*k*_2_, …, *k*_*d−*1_), representing the number of excursions whose diameter is 2, 3, …, *d*−1, with the constraint ∑*k*_*κ*_ = *k*. We write 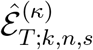 for the ancestries consisting of *k* excursions, all of which have diameter *κ*; and 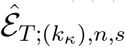 for the set of ancestries that have exactly *k*_*κ*_ excursions with diameter *κ*. Then 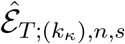 is naturally included in the direct sum of 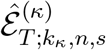. (A sequence of mixed diameters 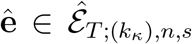 may be decomposed into sequences 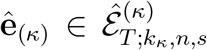 excursions with each particular diameter. Referring back to the example in Figure 4, this would entail making one ancestry by dropping out the green excursions, and a separate one by dropping out the red excursions.) Thus

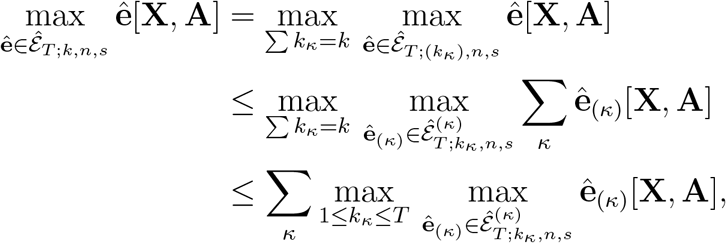

using the general fact that the maximum of a sum is smaller than the sum of maxima. Thus we have

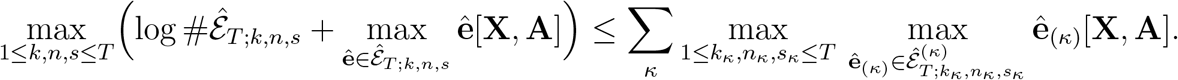

Here *n*_*κ*_ and *s*_*κ*_ simply index the choice of total length and number of change-points among all of the excursions of length *κ*.

Because all excursions in 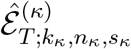 pass through only sites *j* with 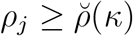, the same argument used for the upper bound in (16) may be applied to show that almost surely for *m >* 0 sufficiently small

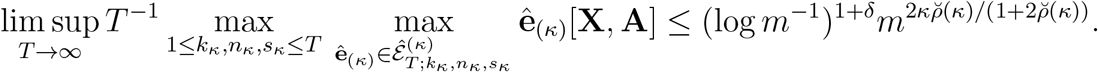

It follows that

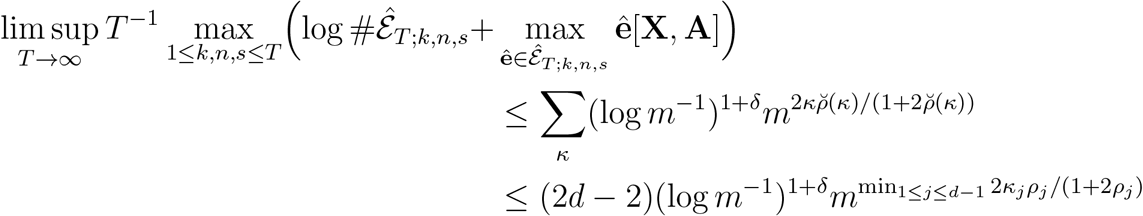

by (17), which completes the proof.

### 5. Derivation of the lower bound

We show that the lower bound applies for each *j*; it will then hold in particular for the *j* at which *κ*_*j*_*ρ*_*j*_ attains its minimum. We may assume without loss of generality that this “second-best” site is *j* = 1, and we will write simply 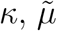, and *<u>ρ</u>* for 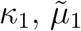, and *<u>ρ</u>*_1_. We assume that the row sums are identically 0, hence there is no cost to migration.

Let 0 = *j*_0_, *j*_1_, *j*_2_, …, *j*_*I*_ = 1, *j*_*I*+1_, …, *j*_*κ*−1_, *j*_*κ*_ = 0, be a cycle from 0 in M, passing through 1. We may fix a real number *A*_*∗*_ and *p >* 0 such that

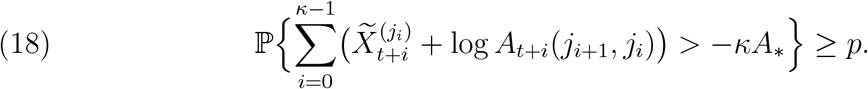

Assume that *m* ≤ e^*−*1^ and *T >* log *m*^*−*1^. Defining 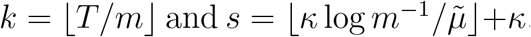, we will apply (12) by considering only excursions of length exactly *s* − 1, which proceed exactly through the sequence of sites 0 = *j*_0_, *j*_1_, …, *j*_*I−*1_, 1, …, 1, *j*_*I*+1_, …, *j*_*κ−*1_, *j*_*κ*_ = 0, where site 1 is repeated exactly *s* − *κ* + 1 times. The idea is that the excursion fills a time block of length *s*, proceeding as quickly as possible from 0 to 1, remaining *s* − *κ* +1 time units at 1, and then returning as quickly as possible to 0.

We define the standard excursion **e**_*◦*_ := (*j*_1_, …, *j*_*I−*1_, 1, …, 1, *j*_*I*+1_, …, *j*_*κ−*1_), with *s* − *κ* + 1 repetitions of site 1; and an ancestry **ê**_*◦*_ consisting of those pairs (*𝓁s* + 1, **e**_*𝓁*_) for which

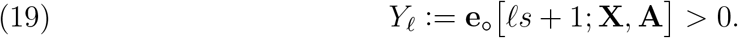

That is, **ê**_*◦*_ is assembled from identical excursions of form **e**_*◦*_ which can start only at times *𝓁s* + 1. Each one of the *k* possible excursions is included precisely when its contribution to the sum would be positive.

We have

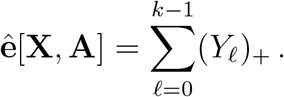

Since the excursion contributions *Y*_*𝓁*_ are all independent, combining (12) with the Strong Law of Large Numbers yields

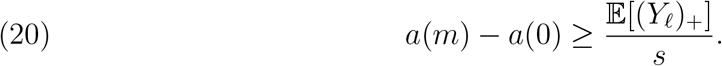

We now observe that for any *𝓁*

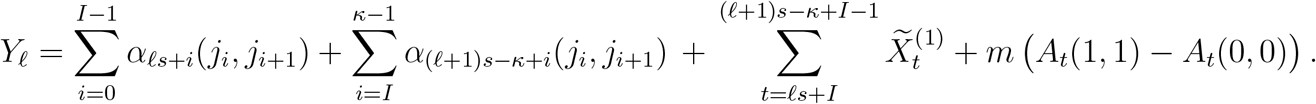

Note that the *α*_*t*_ terms are independent of the 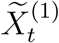 terms, because they draw their component random variables from different times.

By (18) and the definition of *s*, which makes 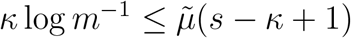, for any *y >* 0

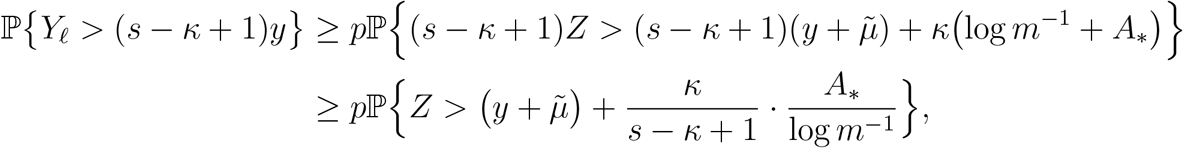

where

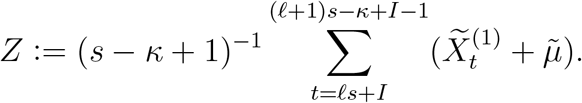

is the mean of independent mean-zero random variables whose moment generating functions have Legendre transform 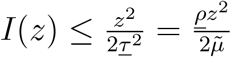. By Cramér’s Theorem [DZ09, Theorem 2.2.3]

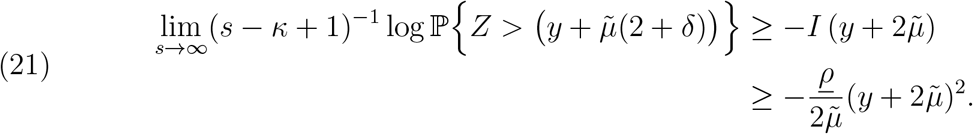

Holding *y* fixed, it follows that for any *δ >* 0 we have for *s* sufficiently large (which is to say, for *m* sufficiently small)

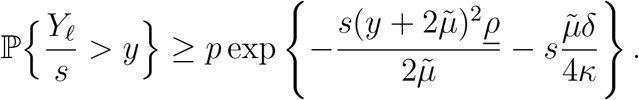

Taking *m* sufficiently small that 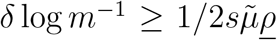 *<u>ρ</u>* and log *m*^*−*1^ ≤ *m*^*−δ/*2^, it follows, by Lemma 4 (proved next), that

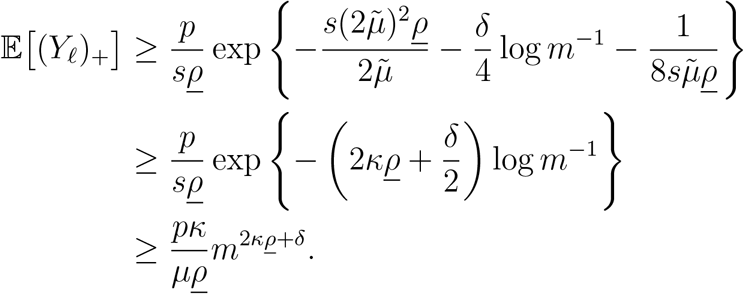

Combining this with (9) and (20) completes the proof.

#### Lemma 4.

*Let Z be a random variable with tail bound*

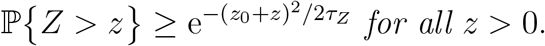

*Then*

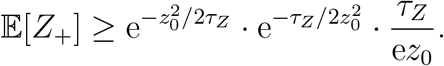

*Proof*.

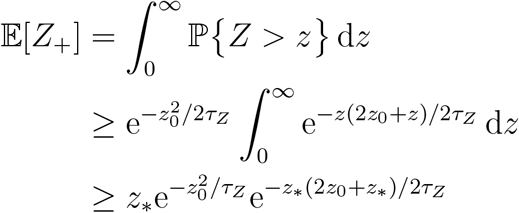

for any fixed *z*_*∗*_ *>* 0, since the integrand is decreasing. Taking *z*_*∗*_ = *τ*_*Z*_*/z*_0_ gives us the result. □

The statements for general *A* follow then from the linear contribution of *m* ∑*A*_*t*_(0, 0) to(20). The rest follows as in the conclusion of section 4 by considering the lower bound, applied to a modified 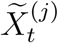, taken as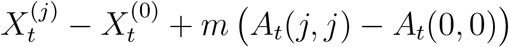, showing that the contribution of *A*(*j, j*) − *A*(0, 0) over all non-trivial excursions can shift the power *η* by an arbitrarily small amount, which may be incorporated into *δ*.

### 6. Proofs of claims

#### 6.1. Derivation of Stirling’s-Formula bound

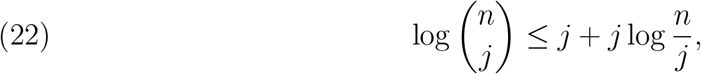

which holds for all positive integers *n* and 0 ≤ *j* ≤ *n*, following the convention 0 · log 0 = 0 · log ∞ = 0. This is because: Since *j* ≥ 0,

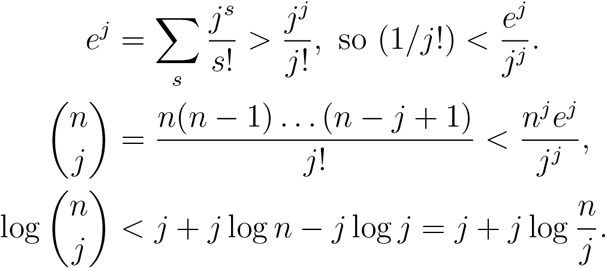

#### 6.2. Proof of Claim 3

*Proof*. We define

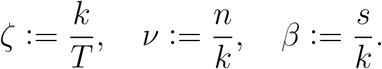

That is, *ζ* is the rate of excursions per unit time; *ν* is the average length of excursions; and *β* is the average number of migration events over the course of the excursions. We have the constraints 1*/ζ* ≥ *ν* ≥ *β* ≥ *κ*_1_ ≥ 2 (since *κ*_1_ is the minimum *κ*_*j*_, hence the minimum number of changes in each excursion). The bound (14) may be written as

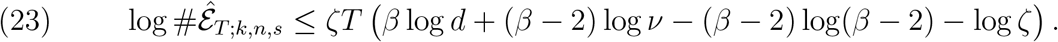

Suppose now we fix some element **ê** of 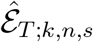 and list all the states of all the excursions in order as *j*_1_, …, *j*_*n*_, and the associated times as *t*_1_ *< t*_2_ *<* · · · *< t*_*n*_. Remember that here *j*_*i*_ = 1 for every *i*. Now, using the definition for an excursion defined above:

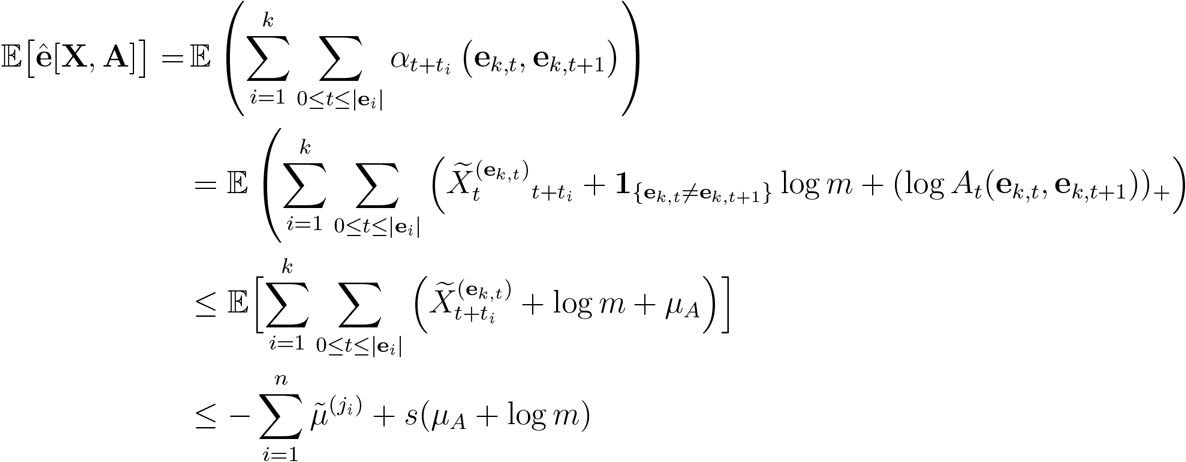

Therefore we have

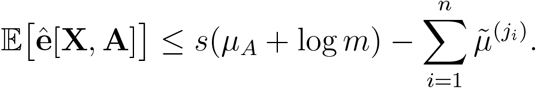

The random variable *Y* := **ê**[**X, A**] − 𝔼[**ê**[**X, A**]] is a sum of sub-Gaussian random variables, hence is also sub-Gaussian. The variance factor is bounded by the sum of the variance factors of the summands, hence no bigger than 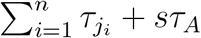. We write

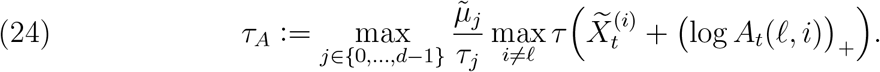

This definition makes the variance factor of 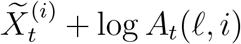 for any *i*≠ 0 will be less than or equal to *τ*_*A*_*/ρ*_*i*_. We are assuming that *ρ*_*j*_ is minimized at so,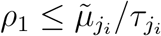 and 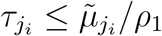 We have

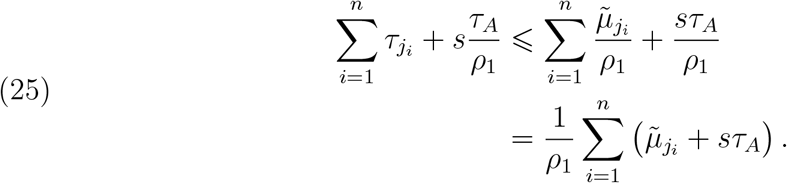

By (3): 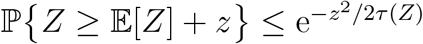 for any *x,z* > 0,

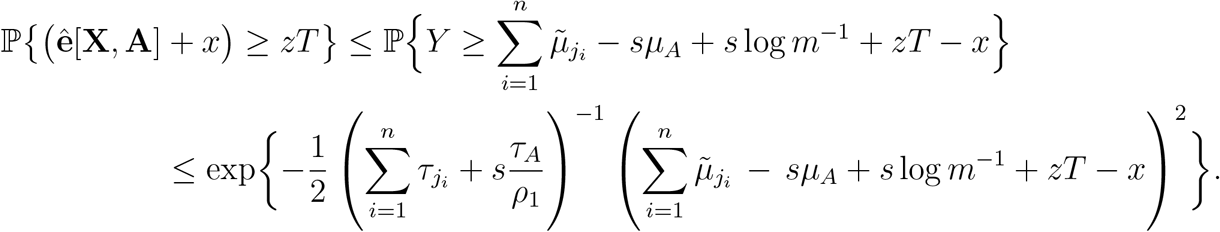

Taking 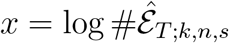 and substituting (23) and (25), we get

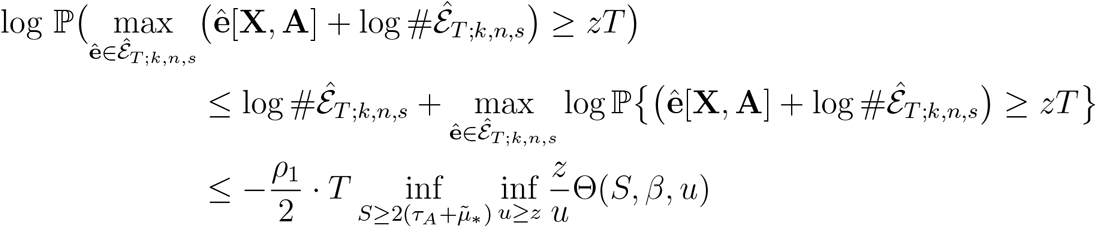

where

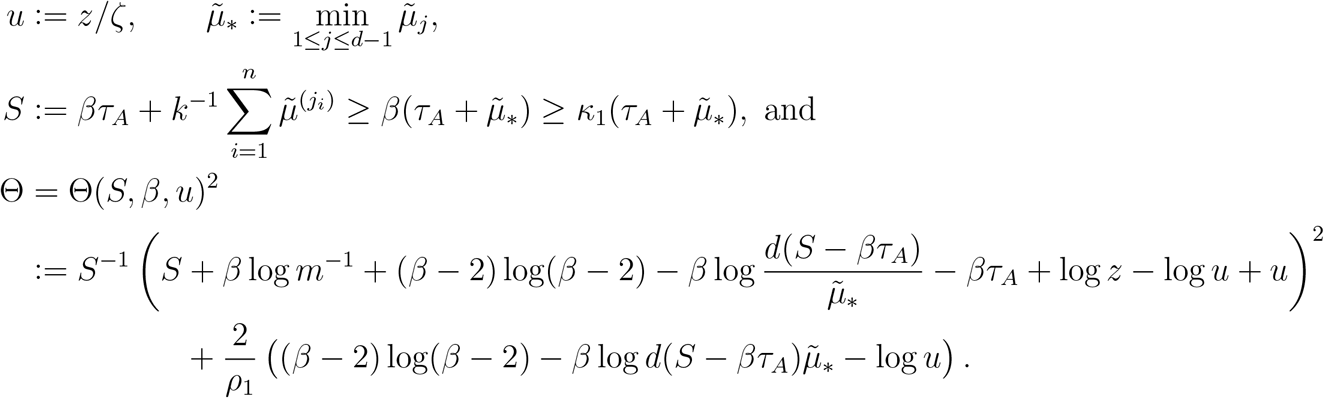

Here we have taken advantage of the fact that log *ζ <* 0, and of the bound 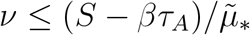.

We will now show that there is a positive constant *m*_0_ — constant in the sense of being expressible in terms only of the model parameters, 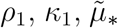 *d, τ*_*A*_, and *µ*_*A*_ — such that for any fixed *m* ∈ (0, *m*_0_) and

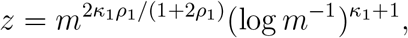

and any *S >* 0, *u >* 0, and *β* ≥ *κ*_1_,

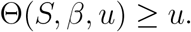

The result then follows immediately, since then for all *T*,

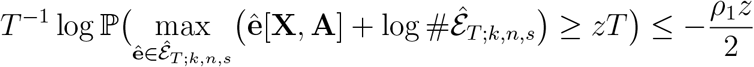

We consider separately the minimization of Θ for large and small values of *S*. Suppose first that *S > *κ**_1_ log *m*^*−*1^. We use the bound *S*^*−*1^(*S* + *x*)^2^ ≥ *S* + 2*x*. We have then

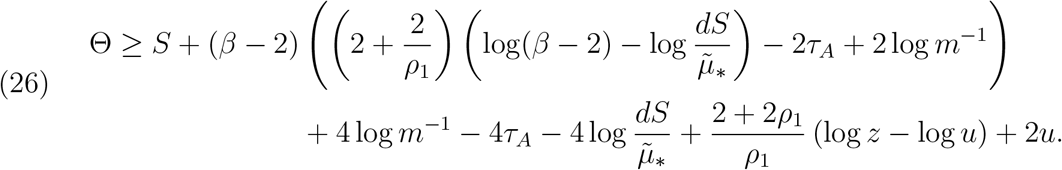

(We have dropped −*βτ*_*A*_ from the log terms, as this would only increase the value of Θ.) By taking the partial derivative with respect to *β*, we see that this is minimized at

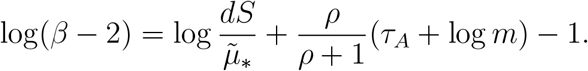

(This value of *β* may be less than the smallest allowable value *κ*_1_, but we may still use it for a lower bound on Θ.) For *m* sufficiently small *S* + log *z >* 0, hence

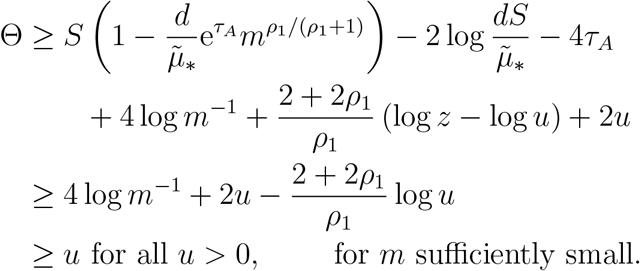

Now suppose that *S* ≤ *κ*_1_ log *m*^*−*1^. We will use the general bound *S*^*−*1^(*S* + *x*)^2^ ≥ 4*x*, which holds for any *S >* 0 and any *y*. We have now a slightly altered version of (26)

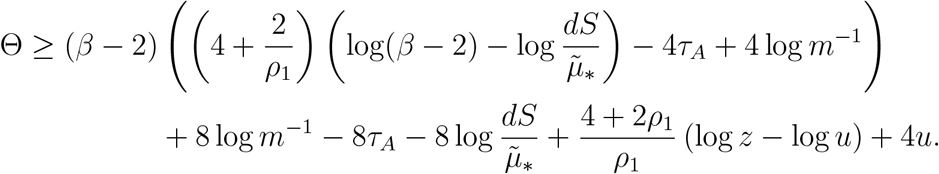

As before, the minimum of this expression with respect to *β* occurs at

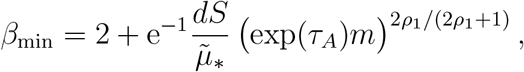

and it is increasing on (*β*_min_, ∞). Since we have assumed *S* ≤ *κ*_1_ log *m*^*−*1^, we may choose the constant *m*_0_ such that this *β*_min_ is strictly less than 3. This means that the minimum will occur at *β* = *β*_min_ when *κ*_1_ = 2, and at *β* = *κ*_1_ otherwise.

Consider first the case *κ*_1_ = 2, so that 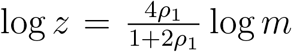. Substituting *β* = *β*_min_ we have

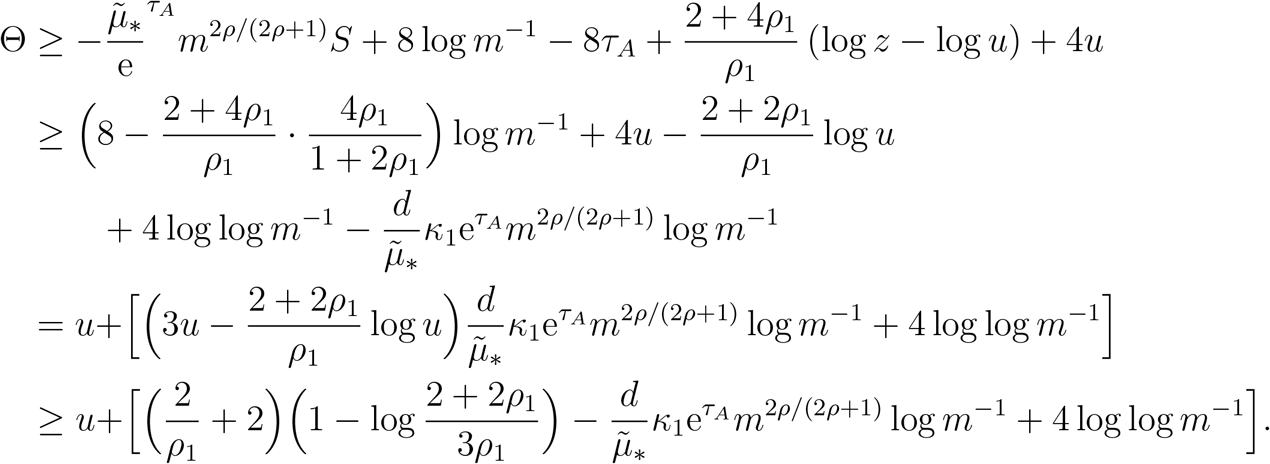

This expression in brackets depends only on model parameters — in particular, not on *u* — and is positive for *m* sufficiently small.

Finally, we consider the case *κ*_1_ ≥ 3. Substituting *β* = *κ*_1_ and simplifying we obtain

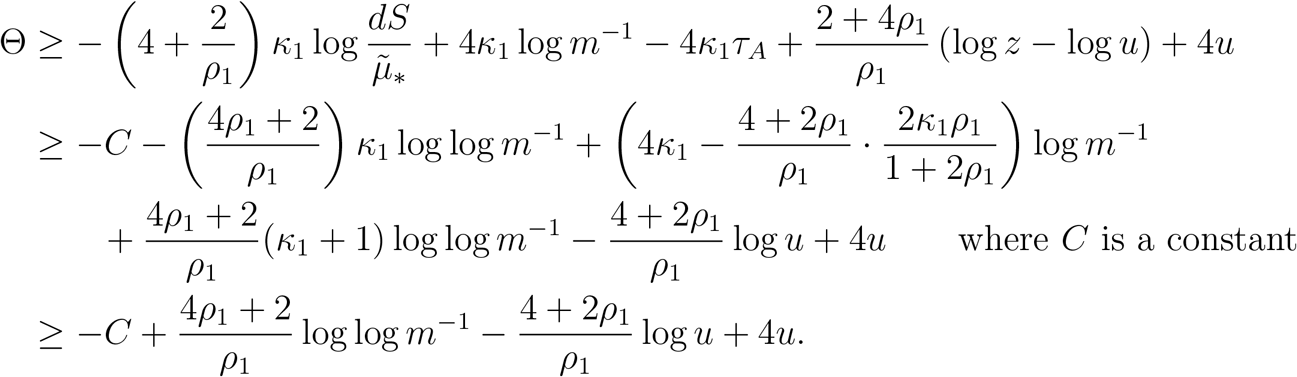

As before, this will be *> u* for all *u >* 0, when *m* is sufficiently small, which completes the proof. □

## Notes

### Competing Interest Statement

The authors have declared no competing interest.

### Summary of Updates

The version reflects comments from reviewers.

